# Cargo-selective regulation of clathrin-mediated endocytosis by AMP-activated protein kinase

**DOI:** 10.1101/2024.11.05.622177

**Authors:** Laura A Orofiamma, Ralph Christian Delos Santos, Ayshin Mehrabi, Nikol Leshchyshyn, Geoffrey G Hesketh, Sadia Rahmani, Aesha Patel, Farnaz Fekri, Rehman Ata, Colin DH Ratcliffe, Mathieu JF Crupi, Lois Mulligan, Morag Park, Anne-Claude Gingras, Costin N Antonescu

## Abstract

The cell surface abundance of many proteins is controlled by clathrin-mediated endocytosis (CME). CME is driven by the assembly of clathrin and other proteins on the inner leaflet of the plasma membrane into clathrin-coated pits (CCPs). Regulation of CCP dynamics allows for control of the function of specific cell surface proteins, impacting a range of cellular outcomes. AMP-activated protein kinase (AMPK) becomes activated upon metabolic insufficiency and facilitates cellular adaptation to nutrient stress. Here, we examined how AMPK regulates CME and the cell surface membrane traffic of β1-integrin. We find that AMPK controls CCP dynamics and regulates the abundance of the endocytic adaptor protein Dab2 within CCPs in a manner that requires the GTPase Arf6, thus selectively promoting the CCP recruitment and internalization of β1-integrin. This reveals a novel signaling pathway for cargo-selective metabolic regulation of CME by AMPK, impacting the function of cell surface proteins such as integrins.

## Introduction

Cells rely on various mechanisms to adapt to fluctuating nutrient availability. An important component of this cellular metabolic regulation is the conserved energy sensor and regulator, AMP-activated protein kinase (AMPK) that becomes activated under various states of nutrient restriction ^1–4^. Understanding how AMPK promotes cellular adaptation in response to fluctuations in nutrient availability is therefore important to understanding metabolic control of cell and systemic physiology.

AMPK responds to a variety of metabolic cues linked to nutrient insufficiency, including an increase in the ratios of AMP:ATP and ADP:ATP, and a reduction of glucose availability for glycolysis. Activation of AMPK promotes metabolic adaptation by stimulating nutrient uptake and catabolic reactions, such as glucose uptake and fatty acid oxidation, and downregulating anabolic processes, such as protein synthesis and cell cycle progression ^5–8^. In addition to a critical role for AMPK activation during severe metabolic challenge, such as in response to ischemia, AMPK is also activated by a range of physiological glucose concentrations ^9,10^. This indicates that outcomes of AMPK activation are not restricted to severe metabolic challenges, suggesting that AMPK has an ongoing role in regulating cellular physiology in a variety of tissues under normal physiological conditions.

One outcome of AMPK activation is regulation of the rate of internalization and/or recycling of specific cell surface proteins ^11,12^. For example, AMPK activation elevates the cell surface levels of the glucose transporters GLUT1 and GLUT4 ^13–16^ and the fatty acid transporter FAT/CD39 ^17^, while decreasing the cell surface abundance of the Na/K ATPase pump ^18^. We previously used selective biotinylation and mass spectrometry to identify that AMPK activation causes broad, but nonetheless selective changes in the abundance of specific cell surface proteins on the plasma membrane ^19^. In this study, we identified that the adhesion receptor β1-integrin (among other integrins) was depleted from the cell surface upon AMPK activation. Regulation of the cell surface levels of such proteins gates their access to substrates such as nutrients or adhesion sites in the extracellular milieu. Hence, the AMPK-dependent regulation of the cell surface proteome allows for a diverse range of responses when sensing metabolic stress cues. However, the mechanism(s) by which AMPK may control the cell surface abundance of proteins, such as β1-integrin, remain unclear.

Clathrin-mediated endocytosis (CME) is the principal mechanism for internalization of cell surface proteins ^20–23^, thus serving to regulate the abundance and hence the function of proteins such as β1-integrin ^24^. CME involves the formation of clathrin-coated pits (CCPs) composed of clathrin, the adaptor protein complex AP2, and other proteins on the inner leaflet of the plasma membrane ^25–31^. The assembly of these proteins within CCPs is coupled to the recruitment of specific membrane proteins destined for internalization (cargo), such as β1-integrin, followed by membrane budding and scission of the cargo-containing vesicle from the cell surface, and finally, its delivery to specific intracellular compartments.

While CME triggers internalization and thus regulates the cell surface abundance of thousands of proteins, this process can also achieve cargo-selective regulation of internalization ^32^. While the heterotetrameric adaptor protein complex AP2 recognizes tyrosine-based (YXXφ) or dileucine-based ([DE]XXXL[LI]) motifs within cytosolic tails of membrane proteins as a cue for CCP recruitment and subsequent internalization ^33–35^, other adaptor proteins broaden the pool of CME cargo ^32,36^. One such adaptor protein, Dab2, is recruited to CCPs via interactions with the α-ear domain of AP2 ^37^; moreover, Dab2 interacts with conserved NPXY/NXXY motifs within the cytosolic tails of specific cargo proteins, such as integrin β-subunit cytoplasmic tails, to promote their recruitment to CCPs and thus their internalization ^24,38–45^.

Cargo-selective regulation of internalization can be achieved by regulating specific adaptor proteins such as Dab2. Whether and how AMPK remodels the cell surface proteome through control of CME, and how this can achieve selective internalization of a subset of cell surface proteins in response to metabolic cues, remains poorly understood. Interestingly, AMPK gates the activity of Arf6, a small GTPase that acts as a key regulator of membrane traffic. Specifically, AMPK may act as a guanyl exchange factor (GEF) for Arf6 ^46^, a phenomenon that appears to also be conserved in yeast, as the AMPK homolog Snf1 acts as a GEF for the Arf6 homolog Arf3p ^47^. Additionally, we previously showed that AMPK activation alters the levels of GTPase activating proteins (GAPs) and GEFs that regulate Arf6 and other Arf proteins on the cell surface ^19^.

Several lines of evidence suggest that Arf6 may regulate clathrin-mediated endocytosis. At the synapse, Arf6 enhances the production of phosphatidylinositol-4,5-bisphosphate that promotes the assembly of CCPs ^48^. Arf6 may also bind directly to the β2-subunit of AP2 and clathrin in a GTP-dependent manner, which is linked to regulation of CCP assembly by Arf6 ^49,50^. Additionally, Arf6 localizes to CCPs ^51^ and promotes the recruitment of AP2 and clathrin to membranes ^52^. Furthermore, GTPase activating proteins such as ARFGAP1 ^50^ or SMAP1 ^53^ that act on Arf6 also interact with CCP components, supporting the model of Arf6 regulation of clathrin endocytosis. In addition to regulating CME, Arf6 is a regulator of clathrin-independent endocytosis ^54–57^ and functions to promote intracellular membrane traffic such as the recycling of integrins, the EGF receptor, and the RET receptor among other cargo proteins ^58–62^. While Arf6 may be involved in a broad range of membrane traffic stages, Arf6 localizes to CCPs and enhances the assembly of specific proteins within CCPs. How Arf6 may function in response to AMPK activation to regulate CCP formation and clathrin-mediated endocytosis of specific cargo proteins remains unexplored.

To resolve how metabolically-derived signals leading to AMPK activation control β1-integrin membrane traffic, we investigated how AMPK regulates CCP composition and dynamics during clathrin-mediated endocytosis. Here, we used total internal reflection fluorescence microscopy (TIRF-M) to analyze clathrin structures at the plasma membrane and show that AMPK activation regulates CCP size and dynamics in a cargo-selective manner. We demonstrate that AMPK activation promotes β1-integrin endocytosis by enhancing Dab2 and Arf6 recruitment to CCPs. Examination of the Arf6 interactome using BioID and other methods reveals a GTP-dependent Arf6 association with clathrin-associated proteins. Taken together, our data suggests a model for AMPK-dependent regulation of the cell surface proteome in a cargo-selective manner through control of CME.

## Results

We previously reported that AMPK activation broadly regulates the cell surface proteome and the endomembrane traffic of specific cell surface proteins ^19^. Our results identified β1-integrin as one of the cell surface proteins selectively depleted from the plasma membrane in response to acute AMPK activation by treatment with the specific allosteric AMPK activator A-769662 ^63^. To determine if conditions that mimic physiological triggers of metabolic stress also activate AMPK (such as conditions that induce a high AMP:ATP ratio), we treated ARPE-19 cells with either A- 769662, oligomycin, or 2-deoxyglucose and first confirmed an increase in AMPK activation by detecting the level of phosphorylated substrate Acetyl-CoA Carboxylase (ACC), a well-established substrate of AMPK (**Figure 1A**). Acute treatment with the metabolic stressors oligomycin or 2-deoxyglucose caused a reduction in the abundance of β1-integrin on the plasma membrane similar to that observed by treatment with A-769662 (**Figure 1B-C**). We previously used siRNA silencing of AMPK α1/2 expression to confirm that the loss of cell surface β1- integrin upon A-769662 treatment is AMPK-dependent ^19^. These data suggest that a range of stimuli that trigger AMPK activation also lead to regulation of β1-integrin membrane traffic.

**Figure 1.**
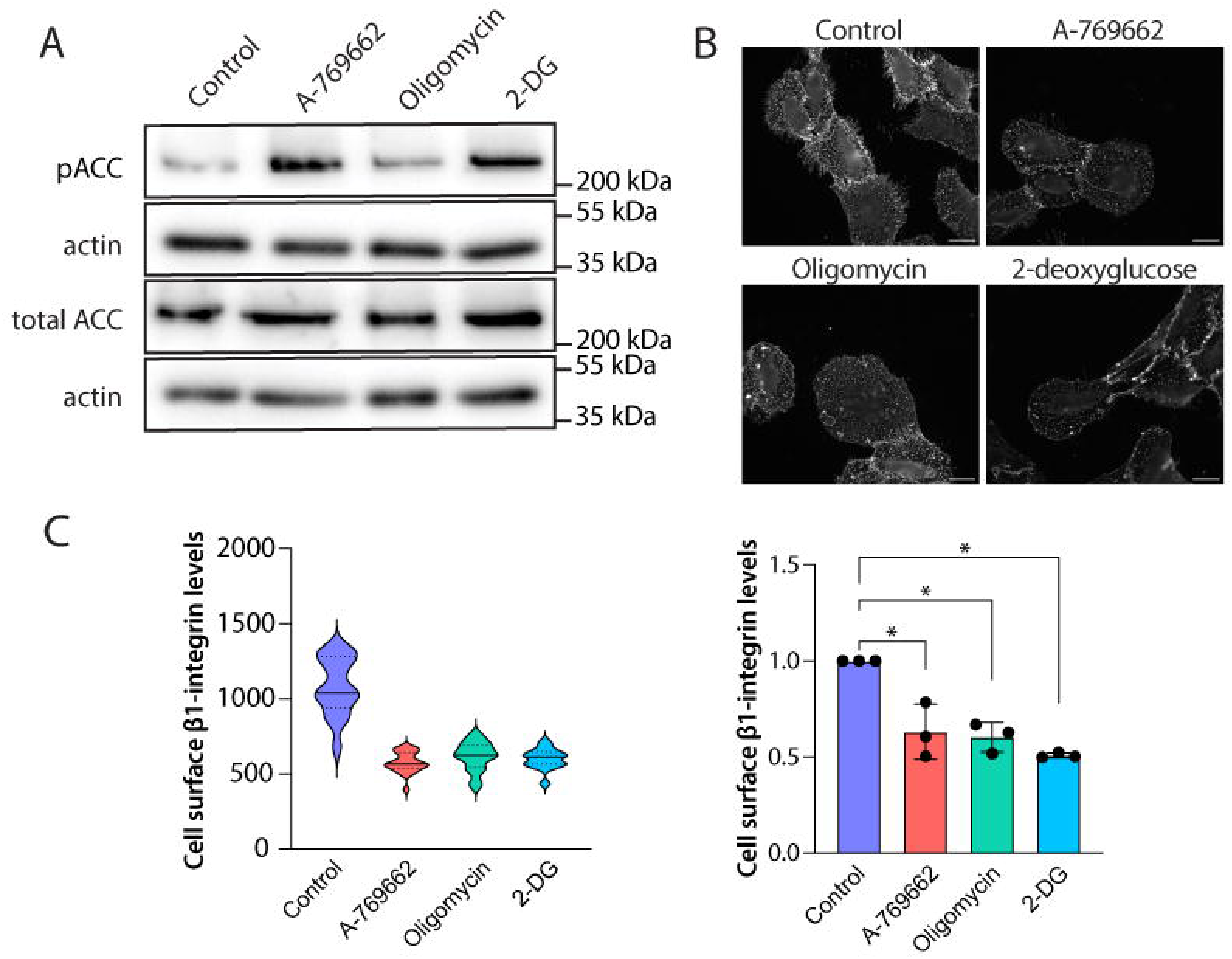
AMPK activation reduces cell surface β1-integrin abundance. (A) Representative immunoblot of ARPE-19 whole cell lysate treated with 100 μM A-769662 for 5 minutes, or 5 μM oligomycin for 20 minutes, or 5 mM 2-deoxyglucose (2-DG) for 20 min, or a vehicle control treatment. (B) Widefield epifluorescence images of fixed ARPE-19 cells treated as in panel A and labelled for cell surface β1-integrin (4B7 antibody clone), as described in *Methods*. Scale bar represents 20 μm. (C) Mean fluorescence intensity of cell surface β1-integrin from a representative experiment (*left*) and shown as the mean ± SD from three independent experiments (*right*) that analyzed 30-50 individual cells per condition. * p<0.05 [one-way ANOVA with Dunnett’s post-hoc test].

### AMPK selectively regulates β1-integrin endocytosis

The reduction of steady-state cell surface β1-integrin in response to acute AMPK activation could be explained by either an enhanced rate of internalization or downregulation of recycling to the plasma membrane. Measurement of β1-integrin recycling up to one hour revealed that AMPK activation does not appreciably impact the rate of β1-integrin recycling (**Figure 2A**). This indicates that AMPK may instead enhance the rate of internalization of β1-integrin.

**Figure 2.**
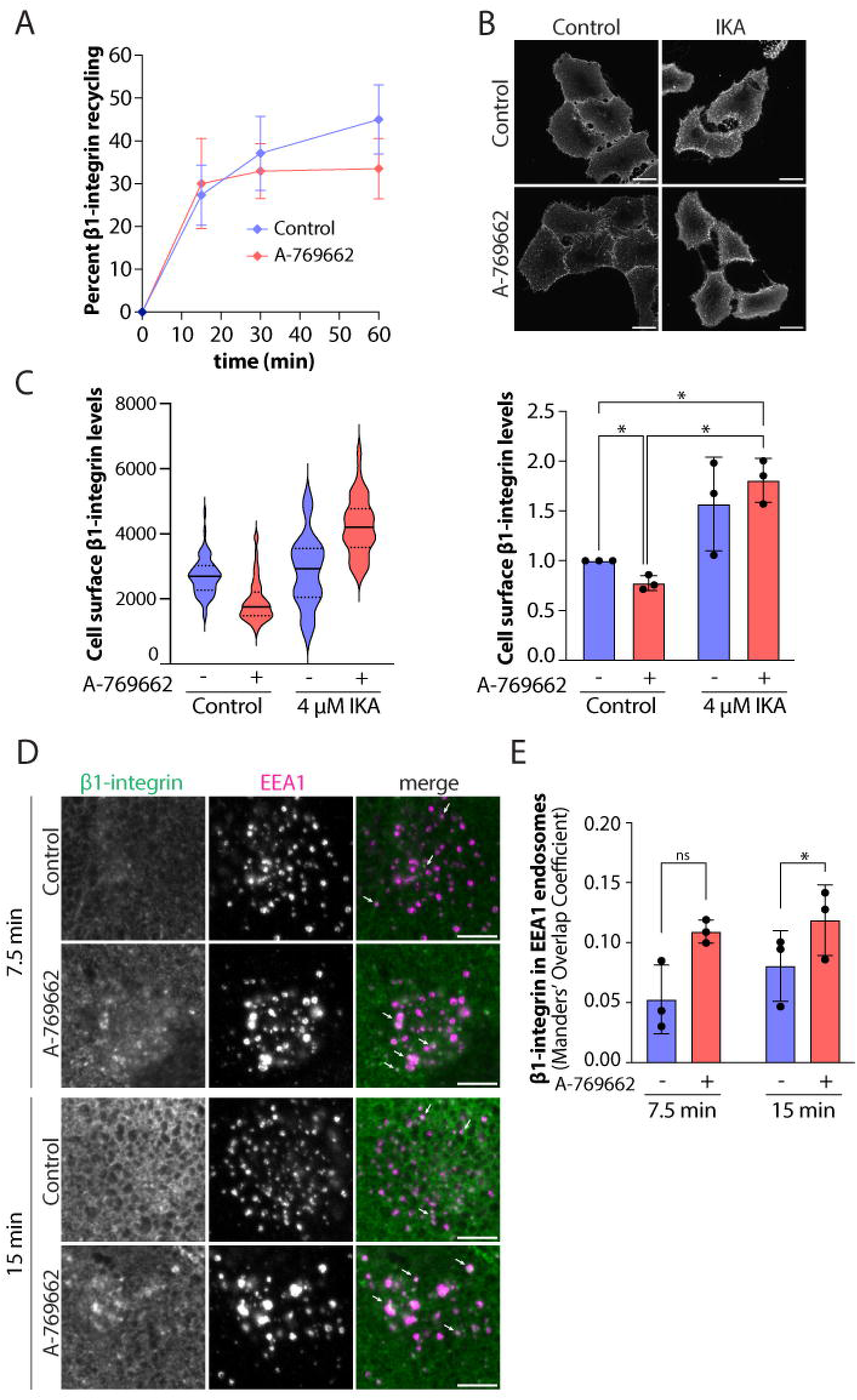
AMPK activation selectively regulates β1-integrin endocytosis. (A) ARPE-19 cells simultaneously treated with 1 mg/mL anti-β1-integrin (antibody clone K20) and either 100 μM A-769662 or vehicle control were fixed at the indicated timepoints. β1-integrin recycling rate shown as a percentage of timed condition fluorescence intensity/total fluorescence intensity. (B) Spinning disk confocal images of fixed ARPE-19 cells pre-treated with 4 μM ikarugamycin (IKA) or vehicle control for 10 minutes, followed by treatment with 100 μM A-769662 or vehicle control for 5 minutes and antibody labelling of cell surface β1-integrin as described in *Methods*. Scale bar represents 20 μm. (C) Mean fluorescence intensity of cell surface β1-integrin from a representative experiment (*left*) and shown as the mean ± SD from three independent experiments (*right*) that analyzed >50 individual cells per condition. * p<0.05 [two-way ANOVA with a Fisher’s LSD test]. (D) ARPE-19 cells simultaneously treated with 1 mg/mL anti-β1-integrin (antibody clone K20) and either 100 μM A-769662 or vehicle control for 15 minutes at 37C. Shown are representative z-series slices of spinning disk confocal images of fixed cells labelled for β1-integrin and early endosome marker EEA1. Scale bar represents 5 μm. (E) Manders’ M1 overlap coefficient for β1-integrin in EEA1-labelled endosomes shown as the mean ± SD from three independent experiments that analyzed 20-30 individual cells per condition. * p<0.05 [two-way ANOVA with a Šídák post-hoc test].

Several distinct routes of internalization have been identified for β1-integrin, generally classified as either clathrin-dependent or clathrin-independent. As clathrin-dependent endocytosis is the major pathway facilitating integrin endocytosis ^64^, we investigated the possibility that AMPK activation may regulate the clathrin-mediated endocytosis of β1-integrin. To test this, we treated ARPE-19 cells with the selective inhibitor of clathrin-mediated endocytosis ikarugamycin ^65^ and measured β1-integrin cell surface abundance in response to AMPK activation. Cells preincubated (10 min) with ikarugamycin exhibited a significant increase in cell surface β1- integrin levels, which was then unaffected by addition of A-769662 (**Figure 2B-C**). Consistent with previous observations, A-769662 treatment alone reduced the level of β1-integrin at the cell surface (**Figure 2B-C**).

To complement these experiments that reveal a loss of cell surface β1-integrin upon AMPK activation with A-769662 treatment, we measured β1-integrin abundance in early endosomes using an antibody-based internalization assay performed in intact cells. Antibodies targeting β1- integrin bind to β1-integrin at the cell surface and accumulate in EEA1-positive early endosomes in a manner proportional to their rate of internalization. We found a significant increase in β1- integrin abundance in EEA1-labelled early endosomes treated with A-769662 compared to control after 15 minutes of internalization (**Figure 2D-E, S1**). Collectively, these data indicate that AMPK activation drives enhanced β1-integrin internalization.

### AMPK exhibits cargo-selective regulation of distinct subsets of CCPs

Regulation of the clathrin-mediated endocytosis of a particular receptor could reflect specific internalization of individual receptors or broader regulation of clathrin-coated pit initiation, assembly, and dynamics. We next examined how AMPK activation may broadly regulate clathrin-mediated endocytosis. To do this, we used TIRF-M to detect clathrin structures and their dynamics on the plasma membrane in ARPE-19 cells stably expressing eGFP-CLCa (henceforth eGFP-CLCa-RPE) and fixed cells at specific times of A-769662 treatment. Acute AMPK activation with A-769662 treatment decreased the fluorescence intensity of clathrin structures, indicating a decrease in the average size of CCPs (**Figure 3A-B**). This suggests that AMPK activation by A-769662 treatment may alter the assembly and/or dynamics of CCPs.

**Figure 3.**
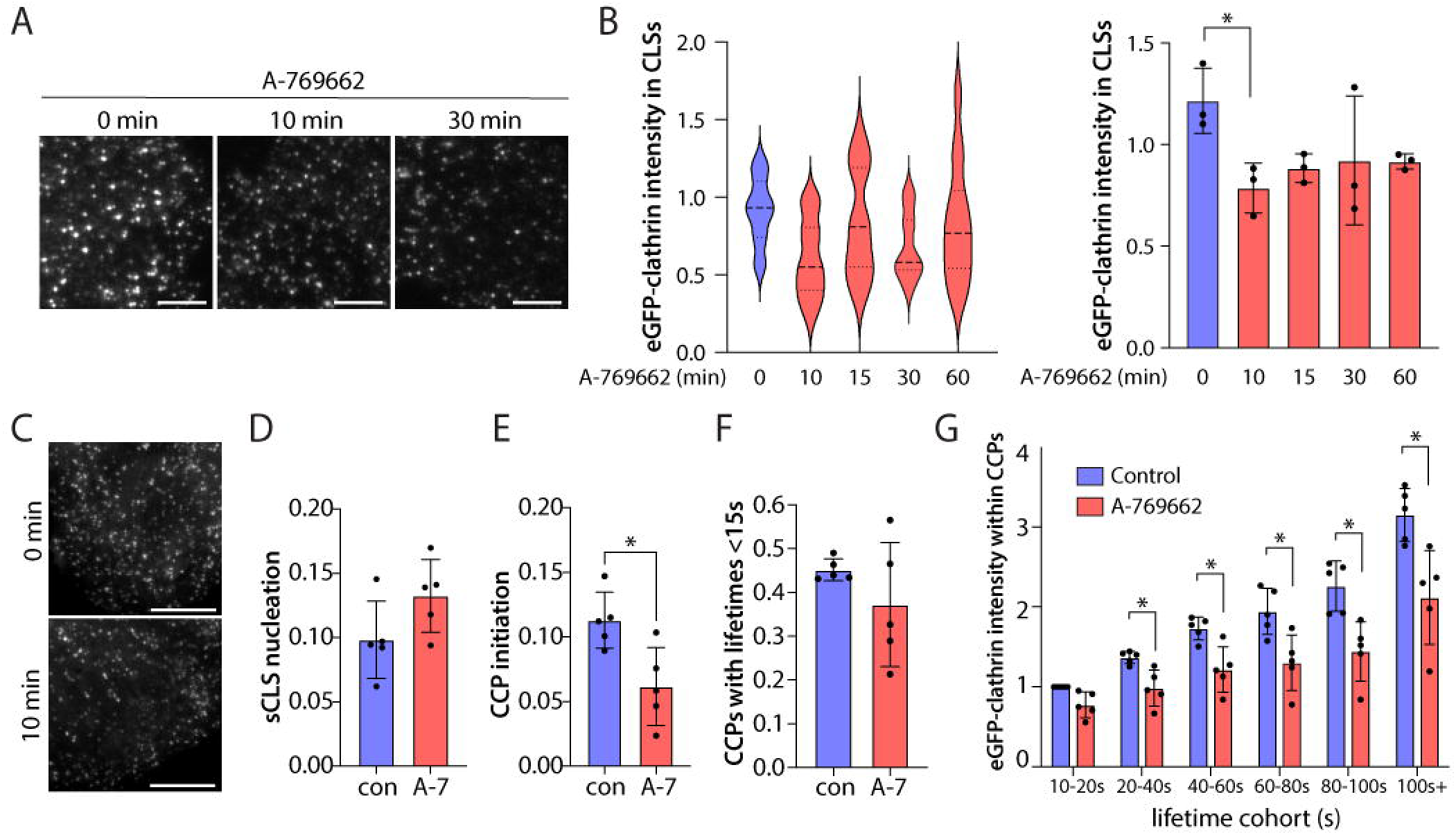
AMPK activation regulates clathrin-coated pit size and dynamics. (A) eGFP-CLCa-RPE cells treated with 100 μM A-769662 or vehicle control were fixed at the indicated timepoints and plasma membrane clathrin structures were visualized with TIRF-M. Scale bar represents 5 μm. (B) Automated detection and analysis of eGFP-clathrin puncta fluorescence intensity reveals CLS size as described in *Methods*. Shown is a representative experiment (*left*) and the mean ± SD from three independent experiments, where for 0 min A-769662: k (cells) = 322 and n (CLSs) = 212852; for 10 min A-769662: k (cells) = 46 and n (CLSs) = 30415; for 15 min A-769662: k (cells) = 46 and n (CLSs) = 33068; for 30 min A-769662: k (cells) = 46 and n (CLSs) = 26592; for 60 min A-769662: k (cells) = 46 and n (CLSs) = 31398. * p<0.05 [one-way ANOVA with Dunnett’s post-hoc test]. (C) Time-lapse TIRF-M imaging of eGFP-CLCa-RPE cells treated with 100 μM A-769662 (or not) were analyzed by automated detection and tracking of CLSs as described in *Methods* to identify bona fide CCPs. Scale bar represents 5 μm. Shown is the mean ± SD from five independent experiments for (D) sCLS nucleation, (E) CCP initiation, (F) CCPs with lifetimes <15 seconds, and (G) eGFP-clathrin intensity within CCPs. Control: k (cells) = 36 and n (CLSs) = 22714; A-769662: k (cells) = 42 and n (CLSs) = 20262. * p<0.05 [Mann-Whitney U test for D-F; two-way ANOVA with a Šídák post-hoc test].

To further resolve how AMPK activation impacts CCP assembly and dynamics, we performed live-cell imaging of eGFP-CLCa-RPE cells coupled to automated detection and analysis of CCPs (**Figure 3C-G, S2A**). This revealed that AMPK activation by A-769662 treatment triggered a decrease in CCP initiation rate (**Figure 3E**), but not in the sub-threshold clathrin-labelled structures (sCLSs) – transient clathrin assemblies that fail to give rise to bona fide CCPs ^33,66^ (**Figure 3D**). This suggests that AMPK activation limits the initial assembly of clathrin and/or other proteins into nascent CCP structures. In contrast to the impact of AMPK on CCP initiation, we did not observe a difference in the proportion of short-lived CCPs upon AMPK activation (**Figure 3F**). Since short-lived clathrin structures include abortive endocytic events that lead to disassembly of CCPs without production of an internalized vesicle, these data suggest that AMPK activation does not alter the stability or likelihood of CCPs developing into longer-lived productive CCPs once they initiate. Consistent with AMPK regulation of the efficacy of initial CCP assembly, the mean intensity of eGFP-clathrin in CCPs in each lifetime cohort also decreased with AMPK activation (**Figure 3G**), similar to the observations of reduced eGFP-CLCa intensity observed in TIRF imaging of fixed samples (**Figure 3A-B**). Changes in eGFP-clathrin in CCPs detected in TIRF-M images could reflect either changes in eGFP-clathrin within individual CCPs or the distance of CCPs from the coverslip (e.g. as a result of alterations of CCP curvature generation). To resolve this, we examined the intensity of eGFP-clathrin in CCPs in time-lapse widefield epifluorescence image series acquired concomitantly to TIRF-M images (**Figure S2B-C**). This revealed that AMPK activation with A-769662 treatment caused a similar reduction of eGFP-clathrin intensity in CCPs in epifluorescence microscopy images as in TIRF microscopy images, indicating that AMPK may cause a reduction in the amount of eGFP-clathrin incorporated within each CCP. These data suggest that AMPK broadly regulates clathrin-mediated endocytosis by controlling initiation and assembly of clathrin structures, while not significantly impacting the stability of the structures that initiate.

We previously reported that AMPK activation triggered a reduction in cell surface β1-integrin levels without affecting the cell surface levels of other proteins such as the transferrin receptor, a well-established cargo of CME ^36^. This suggests that AMPK may selectively regulate the clathrin endocytosis of specific receptors. To investigate this possibility, we measured the internalization rate of the transferrin receptor and the epidermal growth factor receptor using biotinylated conjugates of their ligands ^67^ (Tfn and EGF, respectively) in response to A-769662 treatment. The internalization rates of both transferrin and epidermal growth factor receptors were not significantly impacted in response to AMPK activation by A-769662 treatment (**Figure 4A-B**).

**Figure 4.**
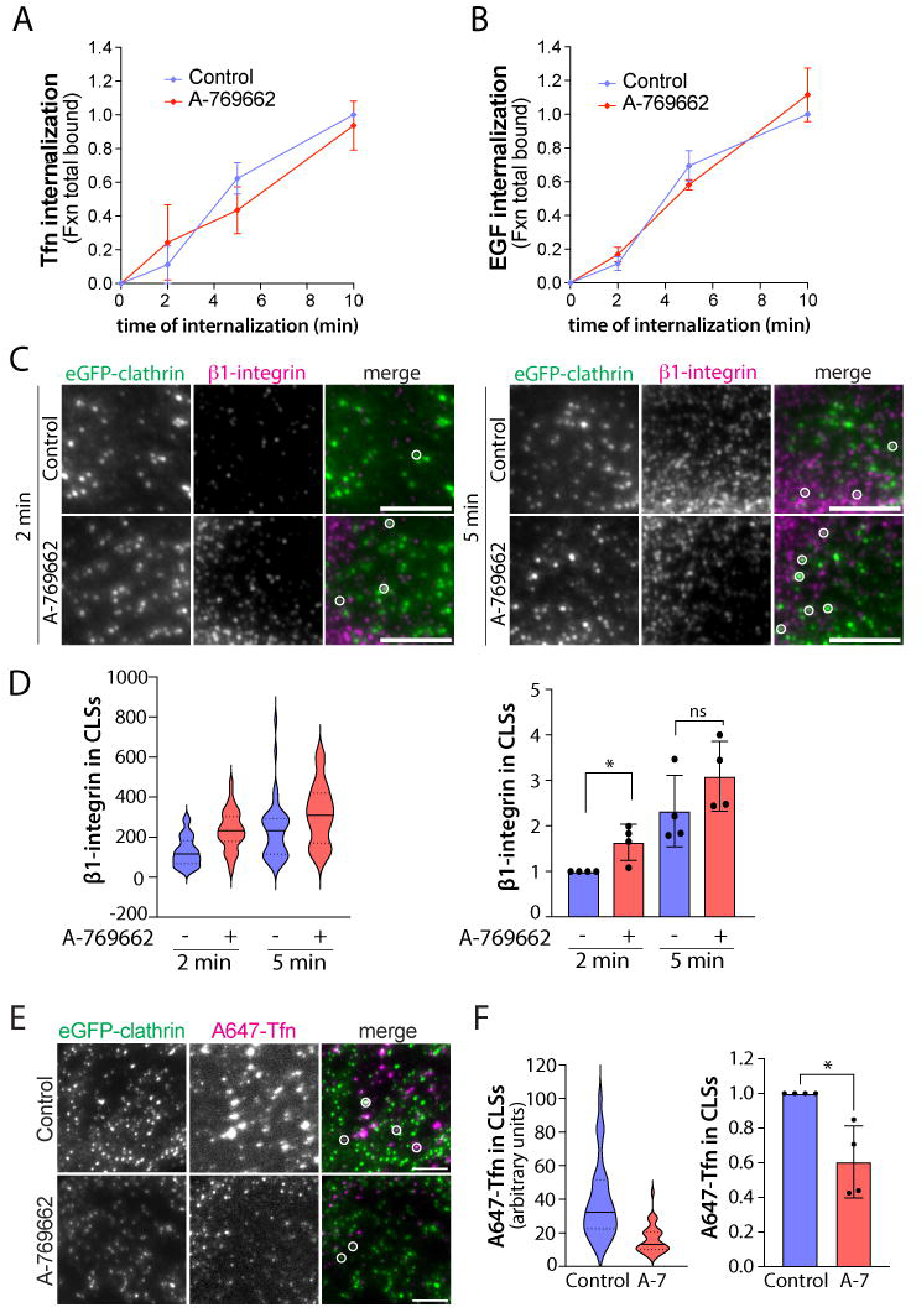
AMPK activation selectively regulates cargo in clathrin structures. ARPE-19 cells treated with either (A) 1 μg/mL biotin-xx-Tfn or (B) 5 ng/mL biotin-xx-EGF in the presence of 100 μM A-769662 or vehicle control treatment were fixed at the indicated timepoints. Tfn and EGF internalization were quantified as described in *Methods*. Shown are the means ± SD from three independent experiments. (C) eGFP-CLCa-RPE cells simultaneously treated with 1 mg/mL anti-β1- integrin (antibody clone K20) and either 100 μM A-769662 or vehicle control at 37C were fixed at the indicated timepoints and imaged by TIRF-M. Scale bar represents 5 μm. (D) Automated detection and analysis of β1-integrin intensity within clathrin structures is shown as a representative experiment (*left*) and as the mean ± SD from four independent experiments (*right*). Control, 2 min: k (cells) = 45 and n (CLSs) = 5708; A-769662, 2 min: k (cells) = 37 and n (CLSs) = 6785; Control, 5 min: k (cells) = 45 and n (CLSs) = 6532; A-769662, 5 min: k (cells) = 49 and n (CLSs) = 7820. * p<0.05 [two-way ANOVA with a Fisher’s LSD test]. (E) eGFP-CLCa-RPE cells were treated simultaneously with A647-Tfn and 100 μM A- 769662 or vehicle control treatment for 5 minutes, fixed, and imaged by TIRF-M. Scale bar represents 5 μm. (F) Automated detection and analysis of clathrin structures and A647-Tfn fluorescence intensity within CLSs are shown as a representative experiment (*left*) and the mean ± SD from four independent experiments (*right*). Control: k (cells) = 34 and n (CLSs) = 7229; A-769662: k (cells) = 37 and n (CLSs) = 8719. * p<0.05 [Mann-Whitney U test].

CCPs are heterogeneous with respect to lifetimes, recruitment of specific cargo receptors, and recruitment of cytosolic endocytic accessory proteins ^28,43,66,68–74^. Indeed, some cargo receptors are recruited to distinct subsets of CCPs that are differentially regulated ^67,68^. Given that AMPK activation by A-769662 treatment results in distinct regulation of β1-integrin internalization compared to that of EGF receptor and Tfn receptor, the regulation of CCPs initiation, dynamics, and size by AMPK could reflect selective regulation of subset(s) of CCPs dependent on cargo therein. To examine this, we exposed eGFP-CLCa-RPE cells to a β1-integrin antibody concomitantly to A-769662 treatment at 37C and measured β1-integrin recruitment to clathrin structures. Acute AMPK activation increased β1-integrin abundance in clathrin structures after a 2 minute incubation period (**Figure 4C-D, S3A**). This suggests that AMPK activation can enhance the recruitment of specific cargo receptors including β1-integrin to CCPs, consistent with the enhanced internalization efficiency of β1-integrin. To complement this approach, we next measured the recruitment of transiently transfected mCherry-β1-integrin to clathrin structures in response to AMPK activation using time-lapse TIRF-M. Consistent with the results of antibody-based detection of β1-integrin, treatment with A-769662 enhanced mCherry-β1- integrin abundance in clathrin structures (**Figure S3B-C**). These data suggest that AMPK activation can enhance the recruitment of specific cargo receptors such as β1-integrin to CCPs, consistent with the enhanced internalization efficiency of β1-integrin.

To resolve if AMPK-dependent regulation of β1-integrin recruitment to CCPs reflected unique regulation of specific cell surface protein cargo during AMPK activation, we measured the abundance of the fluorescent transferrin receptor ligand A647-Tfn in CCPs. In contrast to what we observed for β1-integrin recruitment to CCPs, A-769662 treatment reduced A647-Tfn levels in clathrin structures, indicating that AMPK activation differently impacts the recruitment of various cell surface cargo to CCPs (**Figure 4E-F, S3D**). Collectively, these data suggest that AMPK activation regulates cargo recruitment in a cargo-specific manner, which is likely also coupled to complex regulation of clathrin dynamics leading to enhanced β1-integrin internalization.

### AMPK activation promotes the recruitment of the β1-integrin endocytic adaptor Dab2 to CCPs

The increase in β1-integrin abundance in CCPs suggests that AMPK activation may promote its recruitment to clathrin structures by influencing the presence of cargo-specific adaptor proteins. We considered a possible role for Dab2, an endocytic adaptor protein recruited to CCPs through an interaction with the NPXY/NXXY internalization motif in β1-integrin cytoplasmic tails to facilitate endocytosis ^24^. We therefore investigated whether Dab2 recruitment to CCPs is influenced by acute AMPK activation in eGFP-CLCa-RPE cells using TIRF-M. We found that A- 769662 treatment increased Dab2 abundance in clathrin structures relative to control conditions (**Figure 5A-B, S4A)**.

**Figure 5.**
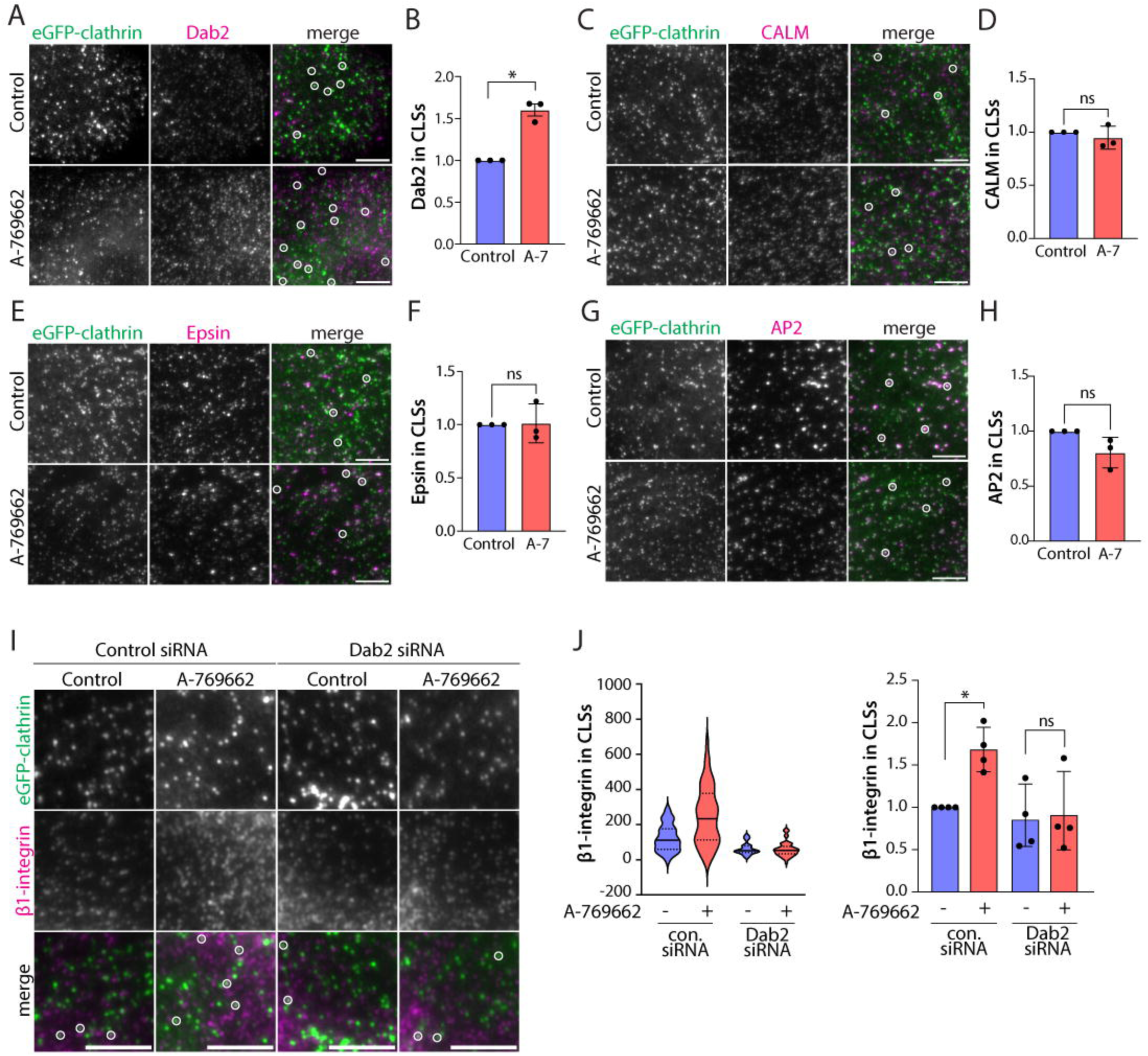
AMPK activation selectively promotes Dab2 recruitment to clathrin structures. eGFP-CLCa-RPE cells treated with 100 μM A-769662 or vehicle control for 5 minutes were fixed and imaged by TIRF-M. Automated detection and analysis of the fluorescence intensity of (A-B) Dab2, (C-D) CALM, (E-F) epsin, and (G-H) AP2 in CLSs were determined by automated detection and analysis of clathrin structures and shown as the means ± SD from three independent experiments. Dab2, control: k (cells) = 147 and n (CLSs) = 47929; Dab2, A-769662: k (cells) = 170 and n (CLSs) = 66109; CALM, control: k (cells) = 38 and n (CLSs) = 15972; CALM, A-769662: k (cells) = 32 and n (CLSs) = 12545; epsin, control: k (cells) = 46 and n (CLSs) = 19642; epsin, A-769662: k (cells) = 39 and n (CLSs) = 17852; AP2, control: k (cells) = 38 and n (CLSs) = 15134; AP2, A-769662: k (cells) = 45 and n (CLSs) = 18130. * p<0.05 [Mann-Whitney U test]. Scale bars represent 5 μm. (I) eGFP-CLCa-RPE cells transfected with Dab2 siRNA or non-targeting control siRNA were simultaneously treated with 1 mg/mL anti-β1-integrin (antibody clone K20) and either 100 μM A-769662 or vehicle control at 37C for 5 minutes. Cells were fixed and imaged by TIRF-M. Scale bar represents 5 μm. (J) Automated detection and analysis of β1- integrin intensity within clathrin structures is shown as a representative experiment (*left*) and as the mean ± SD from four independent experiments (*right*). Control siRNA, control: k (cells) = 49 and n (CLSs) = 9144; Control siRNA, A-769662: k (cells) = 52 and n (CLSs) = 9481; Dab2 siRNA, control: k (cells) = 54 and n (CLSs) = 8193; Dab2 siRNA, A-769662: k (cells) = 49 and n (CLSs) = 7498. * p<0.05 [two-way ANOVA with a Tukey post-hoc test].

The increase in the level of Dab2 in CCPs upon AMPK activation could reflect selective regulation or broad enhancement of protein recruitment to CCPs. To resolve this, we measured the recruitment of other CCP adaptor proteins CALM, epsin, and AP2 in eGFP-CLCa-RPE cells with TIRF-M. Treatment with A-769662 did not change the level of CALM (**Figure 5C-D, S4B**), epsin (**Figure 5E-F, S4C**), or AP2 (**Figure 5G-H, S4D**) recruitment to CCPs. This indicates that AMPK activation leads to selective remodeling of the endocytic accessory proteins within CCPs, with a selective increase in Dab2 recruitment therein.

We next investigated if Dab2 is required to recruit β1-integrin to clathrin structures in response to AMPK activation. To do this, we silenced Dab2 expression in eGFP-CLCa-RPE cells that were then simultaneously exposed to a β1-integrin antibody and A-769662 at 37C. While A- 769662 treatment enhanced β1-integrin recruitment to CCPs in control (non-targeting) siRNA treated cells, the level of β1-integrin in CCPs in Dab2-silenced cells was unaffected by AMPK activation (**Figure 5I-J, S4E-F**). This suggests that β1-integrin is recruited to clathrin structures in a Dab2-dependent manner in response to AMPK activation.

### AMPK-dependent regulation of Dab2 recruitment to CCPs requires Arf6

We next investigated the mechanism by which AMPK may regulate CCP dynamics and composition that can then impact the regulation β1-integrin traffic. Data from our previous report^19^ revealed that AMPK activation controls the plasma membrane abundance of specific Arf GTPase regulators, suggesting the potential regulation of Arf GTPases to effect changes in clathrin endocytosis. Consistent with this, AMPK uses a non-canonical mechanism to enhance the GTP loading of Arf6 ^46^. Additionally, among the mammalian Arf GTPases, Arf6 is most closely associated with the plasma membrane and is a known regulator of β1-integrin traffic ^75,76^. We therefore investigated the possibility that Arf6 is targeted to plasma membrane clathrin structures by AMPK to regulate β1-integrin endocytic membrane traffic.

To determine if Arf6 interacts with β1-integrin and CME components, we performed BioID in HeLa cells transfected with either wild-type Arf6 (Arf6-WT-BirA*), a GTP-bound Arf6 mutant (Arf6-Q67L-BirA*), or a GDP-bound Arf6 mutant (Arf6-T44N-BirA*). Mass spectrometry analysis of biotinylated interactors revealed Arf6-WT-BirA*, Arf6-Q67L-BirA*, and Arf6-T44N-BirA* exhibited interactions with a range of proteins, including proteins that regulate membrane traffic and various integral membrane proteins that undergo dynamic regulation of membrane traffic at the plasma membrane and other compartments (**Figure S5A-B**). Notably, within the set of proteins that regulate membrane traffic, Arf6 demonstrated interactions with many components of clathrin-coated pits (**Figure 6A**, *left panel*), and these largely showed selectivity for Arf6 constructs, with the highest levels of interaction detected within GTP-bound Arf6 (Arf6-Q67L). For example, consistent with a role for Arf6 in regulation of clathrin-mediated endocytosis, Arf6 demonstrated an interaction with three of four subunits of the AP2 complex (AP2M1, AP2B1, AP2A1), the early CCP regulator FCHO2, as well as endocytic accessory proteins and cargo adaptors (CALM – also known as PICALM, Dab2, and others). As expected for interactions dependent on GTP-binding of Arf6, the GDP-bound mutant Arf6-T44N exhibited minimal interactions with clathrin-associated proteins (**Figure 6A**, *left panel*). Interestingly, integral membrane proteins such as solute carriers (SLC proteins, **Figure S5A-B**) and integrins (**Figure 6A**, *right panel*) that are regulated by membrane traffic at the plasma membrane and other compartments also demonstrated interactions with Arf6; however, these interactions largely appeared for all Arf6 constructs. This suggests that interactions of Arf6 with proteins that are putative cargo of membrane traffic is not strictly dependent on Arf6 GTP binding, while interactions of Arf6 with regulators of membrane traffic such as clathrin coat components may be dependent on Arf6 binding to GTP.

**Figure 6.**
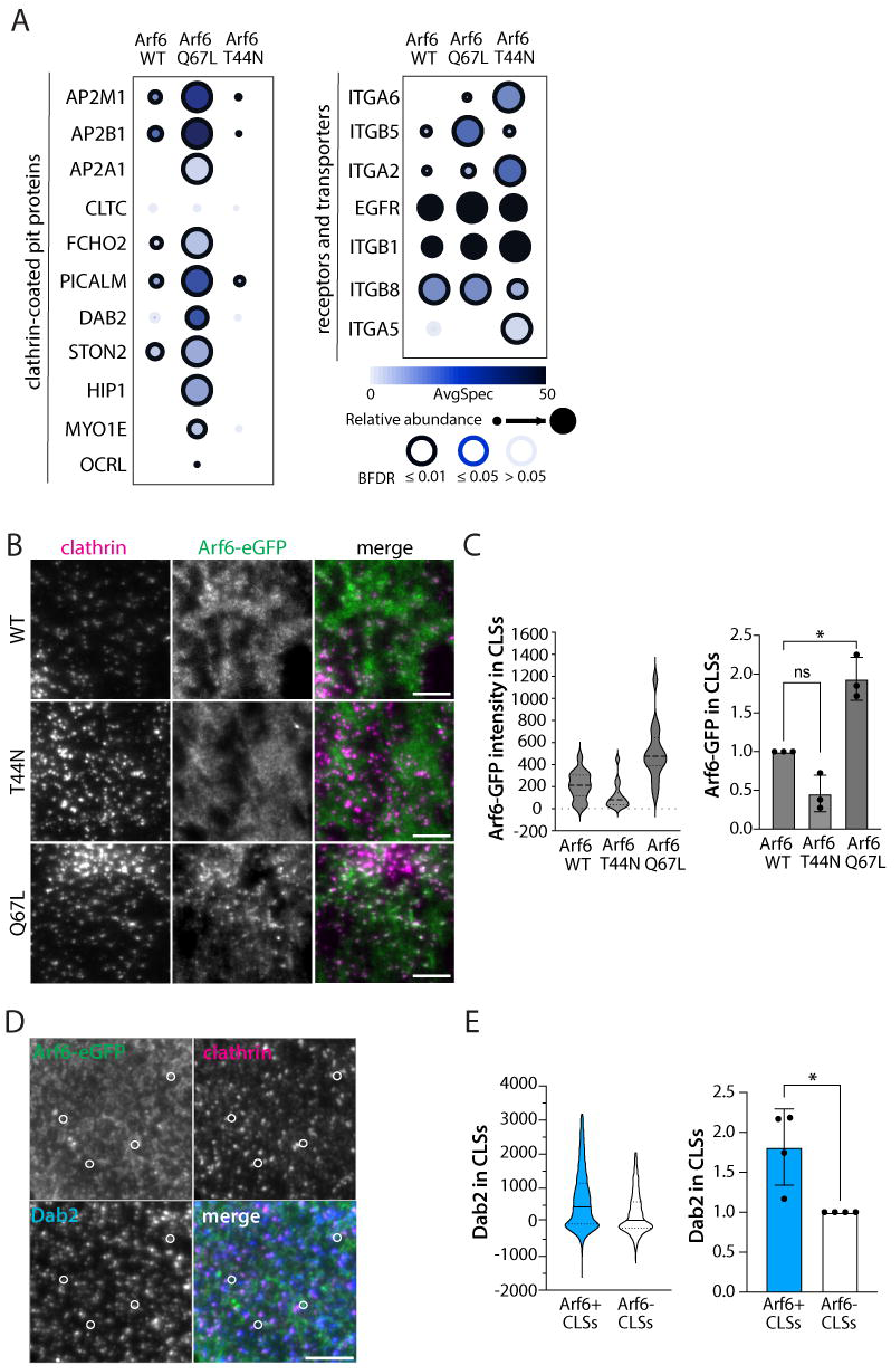
Arf6 associates with clathrin components in a GTP-dependent manner. (A) Dot plot showing prey proteins identified with Arf6-WT-BirA*, Arf6-Q67L-BirA* (GTP-bound), or Arf6-T44N-BirA* (GDP-bound) enriched over endogenous biotinylation (untransfected) and non-specific pan-cellular biotinylation (BirA* alone—BFDR ≤ 1%, SAINT). (B) RFP-CLCa-RPE cells transiently transfected Arf6-WT-eGFP, Arf6-Q67L-eGFP, or Arf6-T44N-eGFP were visualized by TIRF-M. Scale bar represents 5 μm. (C) The fluorescence intensity of Arf6-eGFP constructs in clathrin structures is shown as a representative experiment (*left*) and as the mean ± SD from three independent experiments (*right*). Arf6-WT: k (cells) = 54 and n (CLSs) =20570; Arf6-T44N: k (cells) = 52 and n (CLSs) = 22237; Arf6-Q67L: k (cells) = 54 and n (CLSs) = 25351. * p<0.05 [one-way ANOVA with Dunnett’s post-hoc test]. (D) Arf6-WT-eGFP-RPE cells treated with 1 μM doxycycline for 24 hours were fixed, labelled with anti-Dab2 and anti-clathrin antibodies, and imaged by TIRF-M. Scale bar represents 5 μm. (E) Clathrin structures containing above-threshold Arf6-WT-eGFP intensity levels were determined by automated detection and the fluorescence intensity of Dab2 within structures that either contain Arf6 (Arf6+) or not (Arf6-) are shown as a representative experiment distribution (*left*) and as the mean ± SD from four independent experiments (*right*). k (cells) = 55, n (Arf6+ CLSs) = 10167 and (Arf6-CLSs) = 3694. * p<0.05 [Mann-Whitney U test].

Hence, the differences in the interaction networks observed between the Arf6-Q67L and Arf6-T44N mutants suggests Arf6 may be recruited to CCPs in a GTP-dependent manner. To investigate this, we transfected eGFP-tagged Arf6 cDNA constructs (Arf6-WT-eGFP, Arf6-Q67L-eGFP, and Arf6-T44N-eGFP) into ARPE-19 cells stably expressing RFP-CLCa (RFP- CLCa-RPE) and measured the abundance of each Arf6 construct in clathrin structures using TIRF-M. We found enhanced recruitment of Arf6-Q67L-eGFP to clathrin structures relative to Arf6-WT-eGFP and Arf6-T44N-eGFP (**Figure 6B-C**), while the mean RFP-CLCa intensity in each structure was not significantly different among cells expressing each Arf6 mutant (**Figure S5C-D**). While these images show that Arf6 is recruited to some, but not all CCPs, this further indicates that Arf6 associates with clathrin structures in a GTP-dependent manner.

Under control cell culture conditions, Dab2 has a largely cytosolic distribution and may be limited to recruitment to a subset of clathrin structures at the cell surface ^43^. As both Arf6 and Dab2 appear to be recruited to subsets of CCPs, the detection of proximity between Dab2 and Arf6 by BioID (**Figure 6A**) might suggest some preferential localization of Arf6 to CCPs that also recruit Dab2. We next explored the potential relationship between Arf6 and Dab2 in clathrin structures. To do this, we generated a doxycycline-inducible stable cell line expressing Arf6-WT-eGFP using the sleeping beauty transposon system in ARPE-19 parental cells (Arf6-WT-eGFP- RPE). This system allows tuning of the level of expression to limit overexpression (**Figure S5E**), as we have done previously for other fluorescent protein fusions ^66,74,77,78^. Antibody-based detection of Dab2 and clathrin heavy chain in Arf6-WT-eGFP-RPE cells imaged by TIRF-M allowed for identification and sorting of clathrin-labelled structures based on the presence (Arf6+ CLSs) or absence (Arf6-CLSs) of Arf6-WT-eGFP. We found that Arf6+ clathrin-labelled structures were enriched with Dab2 relative to Arf6-clathrin-labelled structures (**Figure 6D-E, S5F**), which suggests that Arf6 and Dab2 may be recruited to common specific subsets(s) of clathrin structures with unique composition.

To determine if AMPK activation regulates Arf6 recruitment to clathrin structures, we measured the intensity of Arf6-WT-eGFP in clathrin structures using TIRF-M. Acute AMPK activation by A- 769662 treatment significantly increased the level of Arf6-WT-eGFP detected in clathrin structures (**Figure 7A-B, S6A**). To confirm this result, we generated a stable cell line in ARPE- 19 cells encoding an Arf6-WT fusion protein with UltraID – the smallest (<20 kDa) biotin ligase engineered to date that allows for short biotin incubation times to assess acute protein-protein interactions ^79^ (**Figure S6C-E**). Using this system, we treated Arf6-WT-UltraID cells with A- 769662 (or vehicle control) and biotin (or not) for 15 minutes and purified biotinylated interactors using streptavidin pull-down. Immunoblot analysis of the purified biotinylated proteins revealed proximity of AP2 with Arf6, and that A-769662 treatment increased the abundance of biotinylated AP2 when normalized to Arf6-UltraID expression levels (**Figure 7C-D, S6B**). Taken together, these results suggest AMPK activation promotes Arf6 recruitment to clathrin structures.

**Figure 7.**
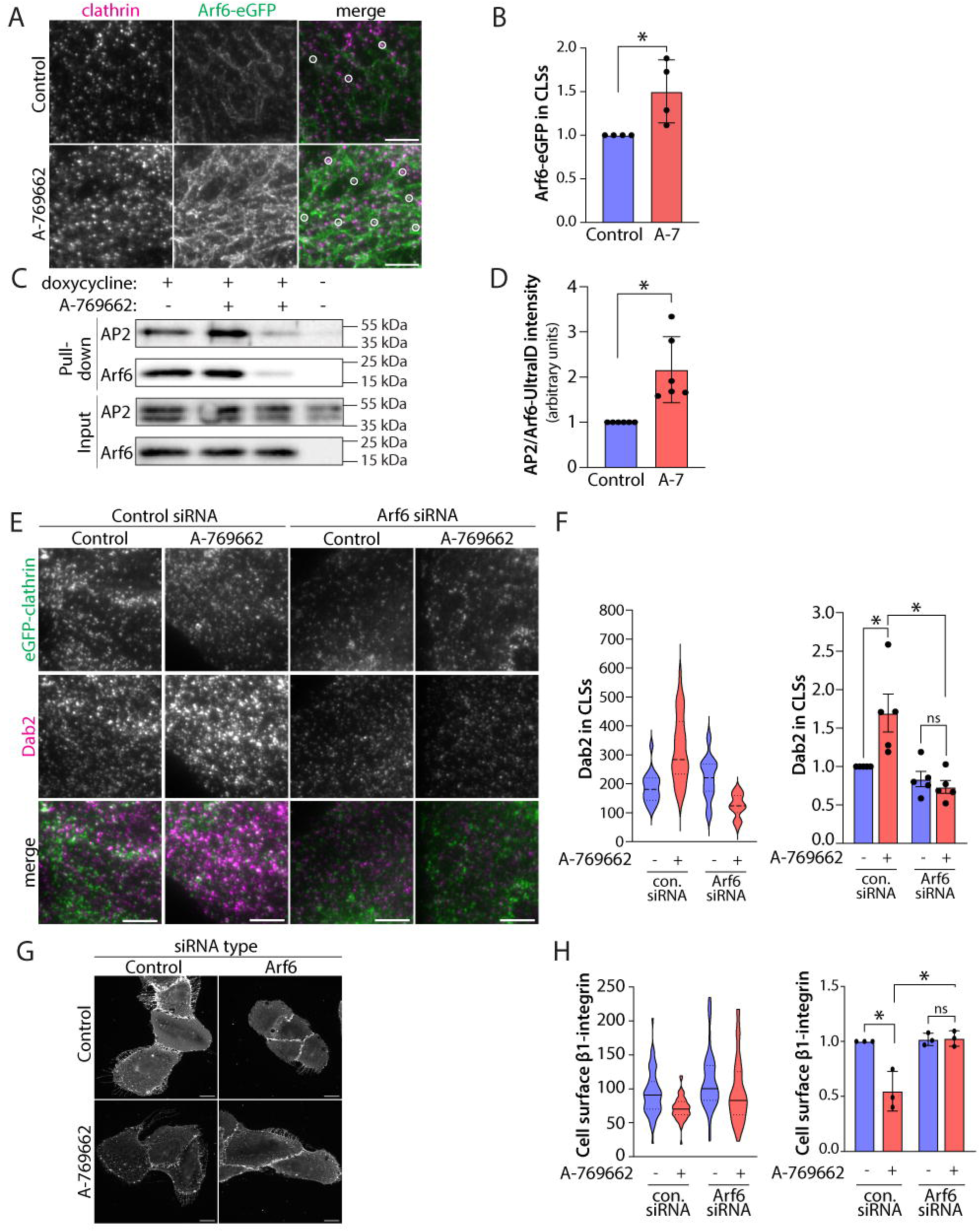
Arf6 is required for AMPK-dependent regulation of Dab2 recruitment to CCPs and β1- integrin internalization. (A) Arf6-WT-eGFP-RPE cells stimulated with 1 μM doxycycline for 24 hours were treated with 100 μM A- 769662 or vehicle control for 5 minutes, fixed, labelled with an anti-clathrin antibody, and imaged by TIRF-M. Scale bar represents 5 μm. (B) Automated detection and analysis of Arf6-WT-eGFP intensity in clathrin structures is shown as the mean ± SD from four independent experiments. Control: k (cells) = 33 and n (CLSs) = 5137; A-769662: k (cells) = 44 and n (CLSs) = 6466. * p<0.05 [Mann-Whitney U test]. The data presented here is a part of the experiment that is also shown in Figure 8A-B and **Figure S6A**. (C) Arf6-WT-UltraID cells stimulated with 1 μM doxycycline (dox) for 24 hours were treated with 50 μM biotin in the presence of 100 μM A-769662 or vehicle control for 15 minutes. Shown is a representative immunoblot of whole cell lysates (input) and biotinylated interactors isolated by streptavidin pulldown (IP) detected by western blotting using the indicated antibodies. (D) Relative abundance of AP2 normalized to Arf6-WT-UltraID expression shown as the mean ± SD from six independent experiments. * p<0.05 [Wilcoxon signed-rank test]. (E) eGFP-CLCa-RPE cells transfected with Arf6 siRNA or non-targeting control siRNA were treated with 100 μM A-769662 or vehicle control for 5 minutes. Shown are TIRF-M images of fixed cells labelled with an anti-Dab2 antibody. Scale bar represents 5 μm. (F) Automated detection and analysis of Dab2 intensity in clathrin structures is shown as a representative experiment (*left*) and as the mean ± SD from five independent experiments (*right*). Control siRNA, control: k (cells) = 177 and n (CLSs) = 74005; control siRNA, A-769662: k (cells) = 166 and n (CLSs) = 76139; Arf6 siRNA, control: k (cells) = 184 and n (CLSs) = 89747; Arf6 siRNA, A-769662: k (cells) = 190 and n (CLSs) = 91736. * p<0.05 [two-way ANOVA with a Šídák post-hoc test]. The data presented here is a part of the experiment that is also shown in Figure 8C-D and **Figure S7C**. (G) ARPE-19 cells transfected with Arf6 siRNA or non-targeting control siRNA were treated with 100 μM A-769662 or vehicle control for 5 minutes and antibody labelled for cell surface β1-integrin (antibody clone 4B7). Scale bar represents 20 μm. (H) Mean fluorescence intensity of cell surface β1-integrin is shown as a representative experiment (*left*) and as the mean ± SD from four independent experiments (*right*) that analyzed 30-50 individual cells per condition. * p<0.05 [two-way ANOVA with a Fisher’s LSD test].

The enhanced recruitment of Arf6 to CCPs upon treatment with the AMPK activator A-769662 and the preferential localization of Arf6 and Dab2 to common clathrin structures suggests that Arf6 could contribute to the elevated Dab2 recruitment to CCPs upon AMPK activation. To determine if Arf6 promotes Dab2 recruitment to CCPs upon AMPK activation, we silenced Arf6 expression using siRNA and measured Dab2 recruitment to plasma membrane clathrin-labelled structures. Silencing Arf6 expression prevented the increase in Dab2 recruitment to clathrin-labelled structures upon acute AMPK activation with A-769662 treatment (**Figure 7E-F, S7A,C**). The requirement for Arf6 to recruit Dab2 to CCPs in A-769662 treated cells could reflect a broad role for Arf6 in regulation of CCP assembly and dynamics, or a specialized role for Arf6 in recruitment of Dab2 to CCPs. To resolve this, we performed live-cell imaging of eGFP-CLCa- RPE cells coupled to automated detection and analysis of CCPs in Arf6 silenced cells and upon treatment with A-769662 (**Figure S8**). Arf6 silencing did not impact CCP initiation (**Figure S8A**) or the proportion of CCPs with lifetimes <15s **(Figure S8B)** in either control or A-769662 treated conditions. In addition, Arf6 siRNA increased the intensity of eGFP-clathrin in some CCPs in both control and A-769662 treated cells (**Figure S8C**). These results suggest that Arf6 modulates some aspects of CCP dynamics but does not appear to play a major role in CCP initiation, initial assembly, or generation of productive, long-lived clathrin structures. This is in turn consistent with a role for Arf6 being limited to regulating the composition of CCPs, such as the recruitment of Dab2 to CCPs.

We next investigated the impact of Arf6 silencing on the cell surface levels of β1-integrin. Consistent with earlier results, A-769662 treatment decreased cell surface β1-integrin levels in cells treated with control (non-targeting) siRNA (**Figure 7G-H**). However, there was no change in cell surface β1-integrin levels in Arf6-silenced cells upon A-769662 treatment (**Figure 7G-H**). We observed similar effects of silencing Arf6 on cell surface β1-integrin levels using a distinct siRNA sequence targeting Arf6 (**Figure S7D-F**), and with a different antibody that recognizes the exofacial epitope of β1-integrin (**Figure S7G-I**). Therefore, Arf6 regulates both Dab2 recruitment to CCPs and cell surface β1-integrin levels in response to acute AMPK activation.

### ArfGAP3 is required for AMPK-dependent regulation of Arf6 and Dab2 recruitment to clathrin coated pits and regulation of integrin traffic

Regulation of GTPases such as Arf6 involves the coordinated action of both GEF and GAP proteins ^80^. While AMPK may act directly as a GEF for Arf6 ^46^, several GAP proteins may regulate Arfs such as Arf6 to allow AMPK-dependent regulation of clathrin endocytosis and β1- integrin membrane traffic. Notably, our previous work that examined the changes in the cell surface proteome of cells upon A-769662 treatment revealed several GAP proteins with GAP activity towards Arf GTPases that change in abundance at the cell surface ^19^, suggesting regulation by AMPK. We thus examined if one of these candidates, ArfGAP3, may contribute to the regulation of clathrin coated pits and integrin membrane traffic following AMPK activation by A-769662 treatment.

To examine how ArfGAP3 may contribute to regulating Arf6 recruitment to plasma membrane clathrin structures, we used cells expressing controlled levels of Arf6-WT-eGFP as described above and TIRF-M. Silencing of ArfGAP3 expression resulted in loss of the A-769662- stimulated gain in Arf6-WT-eGFP within plasma membrane clathrin structures (**Figure 8A-B, S6A, S7B**). Since Arf6 was required for the increase in Dab2 in plasma membrane clathrin structures following AMPK activation with A-769662 treatment, we next examined how ArfGAP3 silencing impacted Dab2 levels within clathrin structures. We observed that silencing ArfGAP3 ablated the increase in Dab2 recruitment to plasma membrane clathrin structures elicited by A- 769662 treatment (**Figure 8C-D, S7C**). Consistent with ArfGAP3 contributing to the gain in clathrin coated pit localization of Arf6 and Dab2 following AMPK activation, silencing of ArfGAP3 also blunted the decrease in cell surface β1-integrin levels following treatment with A-769622 to activate AMPK (**Figure 8E-F**). We observed similar effects of silencing ArfGAP3 on cell surface β1-integrin levels using a distinct siRNA sequence targeting ArfGAP3 (**Figure S7D-F**), and with a different antibody that recognizes the exofacial epitope of β1-integrin (**Figure S7G-I**). Taken together, these results suggest that ArfGAP3 is required to increase the levels of Arf6 and Dab2 within clathrin structures in response to AMPK activation, that in turn promotes the internalization of cell surface β1-integrin.

**Figure 8.**
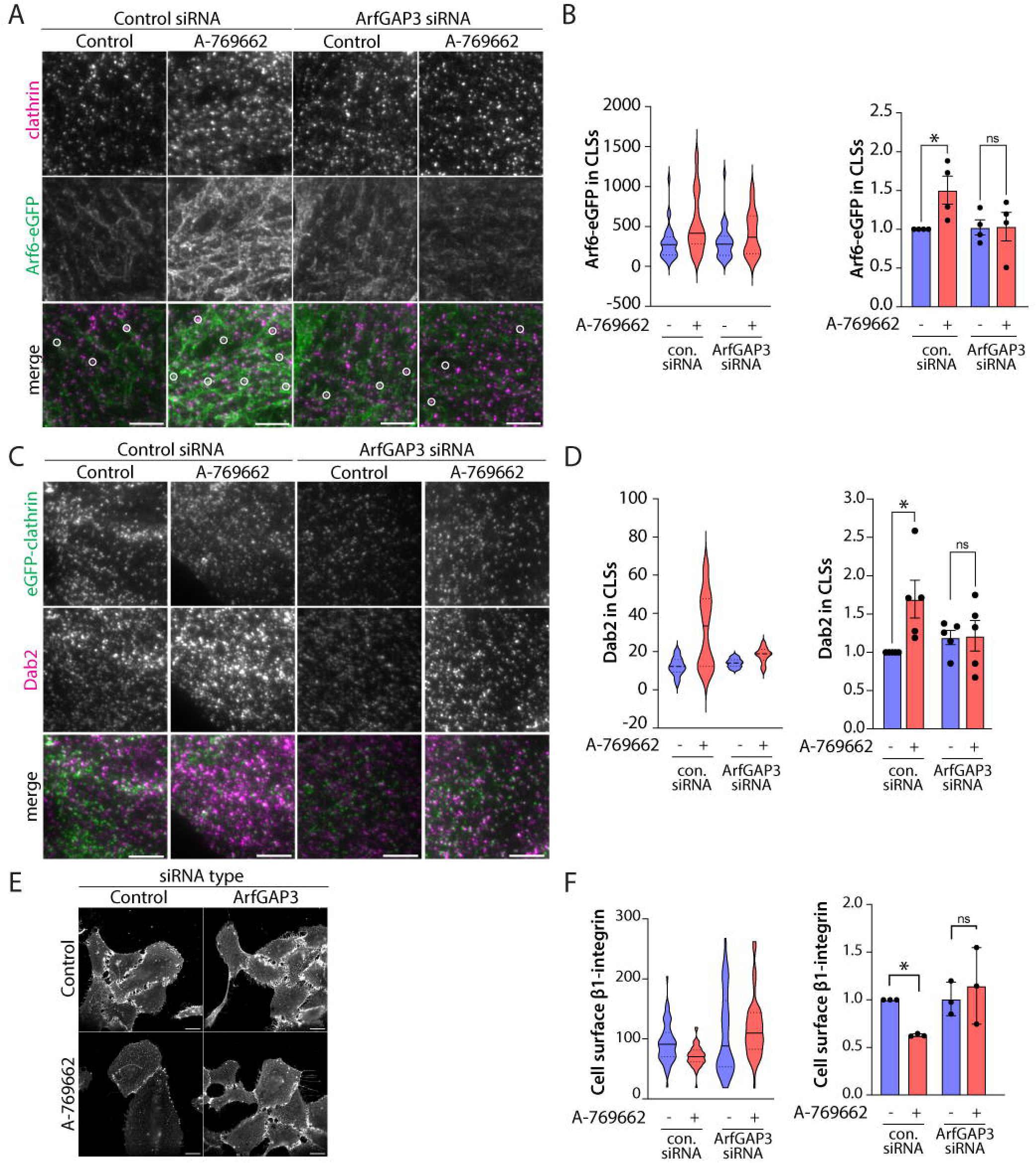
AMPK-dependent regulation of Dab2 and Arf6 recruitment to CCPs requires ArfGAP3. (A) Arf6-WT-eGFP-RPE cells transfected with ArfGAP3 siRNA or non-targeting control siRNA were stimulated with 1 μM doxycycline for 24 hours, followed by treatment with 100 μM A-769662 or vehicle control for 5 minutes. Shown are TIRF-M images of fixed cells labelled with an anti-clathrin antibody. Scale bar represents 5 μm. (B) Automated detection and analysis of Arf6-WT-eGFP intensity in clathrin structures is shown as a representative experiment (*left*) and as the mean ± SD from four independent experiments (*right*). Control siRNA, control: k (cells) = 33 and n (CLSs) = 5137; control siRNA, A-769662: k (cells) = 44 and n (CLSs) = 6466; ArfGAP3 siRNA, control: k (cells) = 32 and n (CLSs) = 5716; ArfGAP3 siRNA, A-769662: k (cells) = 37 and n (CLSs) = 7442. * p<0.05 [two-way ANOVA with a Šídák post-hoc test]. (C) eGFP-CLCa-RPE cells transfected with ArfGAP3 siRNA or non-targeting control siRNA were treated with 100 μM A-769662 or vehicle control for 5 minutes. Shown are TIRF-M images of fixed cells labelled with an anti-Dab2 antibody. Scale bar represents 5 μm. (D) Automated detection and analysis of Dab2 intensity within clathrin structures is shown as a representative experiment (*left*) and as the mean ± SD from five independent experiments (*right*). Control siRNA, control: k (cells) = 177 and n (CLSs) = 74005; control siRNA, A-769662: k (cells) = 166 and n (CLSs) = 76139; ArfGAP3 siRNA, control: k (cells) = 182 and n (CLSs) = 70963; ArfGAP3 siRNA, A-769662: k (cells) = 206 and n (CLSs) = 106193. * p<0.05 [two-way ANOVA with a Šídák post-hoc test]. The data presented here is a part of the experiment that is also shown in Figure 7E-F and **Figure S7C**. (E) ARPE-19 cells transfected with ArfGAP3 siRNA or non-targeting control siRNA were treated with 100 μM A-769662 or vehicle control for 5 minutes and antibody labelled for cell surface β1-integrin (antibody clone 4B7). Scale bar represents 20 μm. (H) Mean fluorescence intensity of cell surface β1-integrin is shown as a representative experiment (*left*) and as the mean ± SD from four independent experiments (*right*) that analyzed 30-50 individual cells per condition. * p<0.05 [two-way ANOVA with a Fisher’s LSD test].

## Discussion

By examining the outcome of AMPK activation on CCP dynamics and composition and on membrane traffic of specific receptors, we reveal selective regulation of clathrin-dependent endocytosis of β1-integrin elicited by AMPK signaling. We find that AMPK activation triggers an increase in Arf6 recruitment to CCPs, which in turn drives increased recruitment of Dab2 and β1-integrin to CCPs, facilitating an increase in internalization. This reveals new insights on mechanisms for cargo-selective regulation of endocytosis and the role of Arf6 in context-specific regulation of clathrin-mediated endocytosis. More broadly, this work provides new insights into how cellular adaptation during metabolic stress may integrate remodeling of cell surface membrane traffic and clathrin endocytosis as key dimensions of AMPK-driven cellular outcomes under these conditions.

### Cargo selective regulation of clathrin coated pits and CME

AMPK activation causes an increase in the internalization of specific cargo such as β1-integrin **(Figure 1B-C, 2B-E, 4C-D)** but does not appreciably affect the rate of internalization of other cargo such as TfR or EGFR **(Figure 4A-B, E-F)**. This suggests that AMPK may distinctly regulate the dynamics and outcomes of different subpopulations of CCPs. We previously showed that TfR and EGFR are largely recruited to different CCPs ^67^. In addition, this previous study revealed that EGFR-positive CCP dynamics are uniquely regulated by PLCγ1-Ca^2+^ signaling that leads to loss of the 5-phosphatase Sjn1 from CCPs, selectively impacting EGFR endocytosis but not that of TfR ^67^. This provides some parallels to the distinct regulation of clathrin-mediated endocytosis of β1-integrin by AMPK activation that we describe here.

Here, we show that the loss of cell surface β1-integrin triggered by AMPK activation is suppressed by the selective ^65^ clathrin endocytosis inhibitor ikarugamycin (**Figure 2B-C**) or by silencing of the clathrin endocytic adaptor Dab2 **(Figure 5I-J)**. Dab2 is an endocytic adaptor for β1-integrin ^24^ and dispensable for TfR endocytosis ^81^. Moreover, silencing subunits of the endocytic adaptor AP2 that are essential for TfR endocytosis does not impact the internalization of Dab2-dependent cargo ^81^. This previous work suggests that clathrin-dependent internalization of Dab2-dependent cargo such as β1-integrin, and other cargo that are strictly AP2-dependent, such as TfR, are largely independent. Our findings in this study are consistent with this, in that we show that AMPK activation leads to increased recruitment of Dab2 and β1-integrin to CCPs along with an increase in β1-integrin endocytosis. In contrast to the increased recruitment of Dab2 and β1-integrin to CCPs, AMPK activation did not significantly change that of AP2, CALM, and epsin (**Figure 5C-H**) within CCPs, and also led to a reduction of TfR recruitment to CCPs **(Figure 4E-F)**. Since AMPK activation does not significantly alter the rate of TfR internalization (**Figure 4A**) or cell surface TfR levels ^19^, decreasing TfR recruitment to CCPs may suggest that AMPK signaling triggers more efficient progression of productive CCPs that harbor transferrin receptor into internalized vesicles. Nonetheless, we observe distinct effects of AMPK on recruitment of β1-integrin and TfR to CCPs that indicates AMPK activation leads to cargo-selective regulation of the clathrin-mediated endocytosis of these receptors.

### Arf GTPase recruitment to, and regulation of, clathrin coated pits

We find that Arf6 is detected within CCPs in a manner that is dependent on Arf6 GTP binding. This is observed both by the detection of Arf6-eGFP in CCPs by fluorescence microscopy, as well as the interactions of Arf6 by proximity biotinylation, both of which suggest a higher degree of interaction of Arf6 with clathrin and clathrin-coated pit proteins in the GTP-bound Arf6 mutant (Arf6-Q67L) compared to the wild-type Arf6 or GDP-bound Arf6 mutant (Arf6-T44N) **(Figure 6A- E, 7A-B)**.

Arf6 could regulate clathrin-mediated endocytosis by broadly enhancing the formation of CCPs or by specifically regulating the function of a specific subset of proteins within clathrin structures. Arf6 binds directly to the β2-subunit of AP2 and clathrin in a GTP-dependent manner and promotes the membrane association of AP2 and assembly of CCPs ^49–52^. In addition, Arf6 GEFs EFA6A, EFA6B and EFA6D interact with Dynamin2 ^82^, and the CCP resident protein endophilin interacts with the Arf6 GEF EFA1 to promote its GTP exchange activity ^83^. These studies suggest that CCPs are able to intrinsically recruit and activate Arf6 therein, in turn suggesting a mechanism for positive feedback with Arf6 GTP loading and CCP membrane assembly that promotes cargo internalization. However, GTPase activating proteins such as ArfGAP1 ^50^ or SMAP1 ^53^ that act on Arf6 also interact with CCP components, supporting the model of Arf6 regulation of clathrin endocytosis. Interestingly, the GAP activity of ArfGAP1 is essential for regulating cargo binding by AP2 ^50^, suggesting that Arf6 has complex functions in control of CCP assembly and cargo recruitment therein.

In contrast, another study found that while Arf6 is recruited to CCPs in an AP2-dependent manner, Arf6 silencing did not impact the internalization of cargo such as TfR ^84^. Instead, Arf6 was required to recruit JNK-interacting protein 3 and 4 (JIP3, JIP4) to CCPs that in turn regulate cargo recycling. This study found that Arf6 silencing did not impact TfR internalization, which is consistent with our study (**Figure 4A**). Moreover, AMPK may directly act in a non-canonical manner as a GEF for Arf6 ^46^, suggesting a potential mechanism for localized Arf6 activation. Our observation that Arf6 silencing impacts AMPK-stimulated β1-integrin internalization (**Figure 7G-H**) suggests that the ability of Arf6 to regulate CCPs dynamics and internalization is restricted to specific subsets of CCPs or to specific physiological contexts.

It remains unclear how Arf6 may regulate CCPs and thus internalization upon AMPK activation. There may be some potential parallels to the regulation of AP1 by Arf1, in which two copies of Arf1 form a bridge to support the conformational change of AP1 from a closed, autoinhibited state to an open conformation capable of interacting with cargo ^85^. The interaction of Arf1 and AP1 that support this conformational regulation are essential for the recruitment of AP1 to the TGN and endosomes. AP2 also undergoes a significant conformational change from a closed to an open conformation, which is enhanced by binding to several molecules of phosphatidylinositol-4,5-bisphosphate (PIP2) ^33,86^ and muniscin proteins ^31^, thus promoting a direct interaction with YXXφ-harbouring cargo proteins for internalization. Hence, it is possible that phosphoinositide-driven regulation of AP2 conformational state predominates for assembly of clathrin structures at the plasma membrane, while Arf-dependent conformational control drives AP1-dependent clathrin structure assembly on the TGN/endosomes.

We show that AMPK activation leads to enhanced recruitment of Arf6 to CCPs, and that Arf6 silencing leads to suppression of the AMPK-dependent increase in Dab2 recruitment to CCPs and loss of cell surface β1-integrin, reflecting β1-integrin internalization. In contrast, AMPK activation did not impact AP2 recruitment therein (**Figure 5G-H**), nor did it impact the internalization rate of TfR or EGFR (**Figure 4A-B**). This further suggests that upon AMPK activation, the regulation of CCP dynamics by Arf6 may not be the direct result of changes in AP2 conformation or CCP localization, but instead to regulate the recruitment of specific endocytic proteins such as Dab2 to CCPs. To support its recruitment to CCPs, Dab2 interacts with the AP2 α-ear domain via a DPF sequence ^45^, and interestingly, the same DPF sequence within Dab2 also supports interaction with the early CCP regulator FCHO2 ^87^.

It remains poorly understood how AMPK activation may promote the enhancement of Dab2 within CCPs in an Arf6-dependent manner. Arf6 promotes the membrane recruitment and activation of PIP5K isoforms that produce PIP2 ^88^. This could suggest that AMPK activation may trigger an Arf6-dependent increase in PIP2 production. In addition to the regulation of AP2 conformation, PIP2 also regulates FCHO2 membrane organization and cargo recruitment ^89^, and directly binds to Dab2 to promote its membrane recruitment ^41,90^. Given the complexity of Arf6 regulation of PIP5K, PIP2 dynamics, and selective cargo recruitment, resolving the contribution of regulation of PIP2 dynamics upon AMPK activation to cargo-selective clathrin endocytosis is beyond the scope of this study, but will be an important priority for future research.

### Regulation of Dab2, Arf6, and integrin traffic by ArfGAP3

We find that AMPK activation triggers changes in Dab2 recruitment to CCPs as well as β1- integrin internalization. We also show that the increase in Dab2 in CCPs triggered by AMPK activation not only requires Arf6, but also the Arf GAP ArfGAP3. Moreover, silencing ArfGAP3 also suppressed the recruitment of Arf6 to CCPs (**Figure 8A-B**) as well as the loss of cell surface β1-integrin stimulated by AMPK activation (**Figure 8E-F**). Hence, these data suggest a model in which AMPK activation may not only trigger an increase in Arf6 GTP loading to regulate the clathrin endocytosis of specific cargo, but may also depend on specific GAPs for regulation of Arf6-dependent membrane traffic outcomes during conditions of metabolic stress.

While ArfGAP3 regulates COPI-mediated traffic ^91,92^, ArfGAP3 has other emerging functions. For example, ArfGAP3 was identified in a screen of Arf GAPs that regulate post-Golgi traffic ^93^, and thus regulates the endomembrane traffic of CIMPR and EGFR. ArfGAP3 also regulates the post-Golgi membrane traffic of the facilitative glucose transporter GLUT4 in myoblasts ^94^. These studies indicate that ArfGAP3 has functions outside of regulation of COPI-dependent membrane transport. Further to this and consistent with our results, ArfGAP3 was identified in a screen for proteins that impact internalization of αvβ6 integrins in a clathrin-dependent manner ^95^. While ArfGAP3 has significant perinuclear localization consistent with Golgi recruitment, clear ArfGAP3 vesicular or punctate structures are also observed throughout the cell ^96,97^. Interestingly, the GAP activity of ArfGAP3 is enhanced by PIP2 ^91^, further strengthening a model in which some of the functions of ArfGAP3 extend beyond its control of COPI-transport. Nonetheless, we are not aware of any study that demonstrated GAP activity of ArfGAP3 towards Arf6 specifically, thus suggesting that the regulation of Arf6 and Dab2 localization by ArfGAP3 may not be direct and could involve complex other factors. Our results add to a growing role for ArfGAP3 in the regulation of membrane traffic outside of control of COPI vesicle formation. Moreover, we reveal that the impact of ArfGAP3 silencing appears to be restricted to the AMPK-activated state, particularly in the regulation of cell surface β1-integrin, suggesting that ArfGAP3 is a conditional regulator of clathrin-dependent internalization and/or other membrane traffic phenomena.

Our results show that ArfGAP3 silencing leads to the loss of AMPK-stimulated enhancement of Arf6 and Dab2 within CCPs (**Figure 8A-D**). It is possible that ArfGAP3 regulates β1-integrin internalization in a manner that requires GAP activity towards Arf6, perhaps by maintaining a pool of Arf6 that can be readily activated by AMPK to thus engage with regulation of the endocytic machinery. Other Arf GAPs including ArfGAP1 ^50^ or SMAP1 ^53^ also function to promote internalization, suggesting that fine control of Arf GTP loading and hydrolysis may be a common and essential dimension of localized regulation of Arf6 control of clathrin-mediated endocytosis. While outside of the scope of the current study, resolving how Arf6 and its GAPs control formation of CCPs will be very interesting to pursue in future studies. Our work nonetheless identifies specific cell physiological states (nutrient starvation leading to AMPK activation) and internalized cargo (β1-integrin) that depend on Arf6 and Arf GAP proteins for clathrin-dependent endocytosis.

### AMPK regulation of integrin endocytosis and membrane traffic

The ability of AMPK activation to regulate CCP dynamics and internalization of selective cargo such as β1-integrin provides another dimension to the regulation of endocytic membrane traffic by metabolic signals, which we reviewed in ^11,12^. While we show regulation of β1-integrin membrane traffic and internalization by AMPK activation, we did not examine if AMPK activation regulates specific β1-integrin heterodimers, defined by interaction with specific alpha-subunits. It will be of interest for future studies to examine if AMPK regulates specific integrin heterodimers, given the unique functions and regulation of these complexes. However, since the NPXY endocytic motif that interacts with Dab2 is present within β1-integrin ^38^, it is plausible that AMPK-dependent regulation of Dab2 recruitment to CCPs may broadly impact a range of β1-integrin heterodimers.

We also reported that the modification of proteins with O-linked β-GlcNAc (O-GlcNAc), which is sensitive to nutrient availability such as that of glucose flux into the hexosamine pathway, regulates CCP size and composition ^74^. This suggests that both O-GlcNAc protein modification and AMPK activation regulate CCP dynamics. While AMPK is also able to regulate the single enzyme required for O-GlcNAc protein modification, O-GlcNAc Transferase (OGT), it is very likely that O-GlcNAc protein modification and acute AMPK activation regulate CCPs in distinct and orthogonal manners. This is due to the fact that we see changes in CCP dynamics and cargo internalization upon acute (5-10 min) activation of AMPK in this study, which are not consistent with the much longer time (2-4 h) of inhibition of OGT required to effect changes in O-GlcNAc protein modification and clathrin endocytosis ^74^.

We report that AMPK activation triggers selective regulation of β1-integrin recruitment to CCPs and β1 internalization, while this does not impact the internalization of TfR or EGFR. While the outcome of AMPK activation is thus cargo selective, the regulation of cell surface membrane traffic by AMPK signaling may extend beyond that of β1-integrin, as we found broad changes in cell surface abundance of many other proteins by AMPK activation ^19^. It is notable that the Arf6 proximity biotinylation interactome reveals a wide range of transport proteins (**Figure S5A-B**). Further, Dab2 is an endocytic adaptor for other proteins such as Low Density Lipoprotein Receptor (LDLR) ^81^, cystic fibrosis transmembrane conductance regulator (CFTR) ^44,98^, and Megalin ^99,100^. It is thus possible that AMPK activation allows broad remodeling of the cell surface proteome by gating the clathrin-mediated endocytosis of a subset of cell surface proteins to allow adaptation to conditions of nutrient scarcity. The extent to which AMPK-dependent regulation of Arf6 and Dab2 that we identify here could impact cell surface proteins beyond β1-integrin is worthy of investigation.

In conclusion, we find that acute AMPK activation selectively regulates a subset of clathrin structures to control the internalization of specific cell surface cargo proteins such as β1- integrin. Activation of AMPK enhances Dab2 recruitment to CCPs in a manner dependent on Arf6 GTP-binding and the presence of ArfGAP3. The extent to which the loss of cell surface β1- integrin in response to acute activation of AMPK impacts integrin-dependent cell survival, adhesion, and migration remains an important area of investigation. Our results emphasize the role of metabolic stress cues and nutrient availability in remodeling the cell surface proteome through clathrin-mediated endocytosis.

## Data and Resource Availability

Requests for further information and resources should be directed to and will be fulfilled by the lead contact, Costin Antonescu. Plasmids generated in this study will be deposited in Addgene upon publication. All unique/stable reagents generated in this study are available from the lead contact with a completed materials transfer agreement.

## Acknowledgements

This work was supported by Discovery Grants from the Natural Sciences and Engineering Research Council (NSERC) to C.N.A. (RGPIN-2016-04371) and a Foundation operating grant to A.C.G. (143301) from the Canadian Institutes of Health Research (CIHR). L.A.O. was supported by a Doctoral Research Award and a Canada Graduate Scholarship – Masters from CHIR as well as an Ontario Graduate Scholarship (OGS). Proteomics work was performed at the Network Biology Collaborative Centre at the Lunenfeld-Tanenbaum Research Institute, a facility supported by Canada Foundation for Innovation funding, by the Ontario Government, and by Genome Canada and Ontario Genomics (OGI-139).

## Declaration of Interests

The authors declare no competing interests.

## Supplemental information

Document S1: Figures S1-S8 and Tables S1-2 Table S3

## Methods

### Cell culture

Human wild-type retinal pigment epithelial cells (ARPE-19, male, herein RPE-WT; obtained from American Tissue Type Collection (ATCC, Manassas, VA) and ARPE-19 cells stably expressing clathrin light chain fused to enhanced GFP (eGFP-CLCa-RPE) ^66,67,72,101^ were cultured in Dulbecco’s Modified Eagle Medium/Nutrient Mixture F-12 (DMEM/F-12; Gibco) supplemented with 10% fetal bovine serum (FBS; Gibco), 100 U/mL penicillin and 100 μg/mL streptomycin (P/S; Gibco) at 37C and 5% CO_2_. Cells were passaged upon reaching ∼80% confluency and lifted with 0.25% Trypsin/EDTA (Gibco).

### Stable transfections using Sleeping Beauty transposon system

pSBtet-BP was a gift from Eric Kowarz (Goethe-University of Frankfurt, Frankfurt, Germany, plasmid 60496; Addgene; http://n2t.net/addgene:60496; RRID:Addgene_60496) ^102^. pCMV(CAT)T7-SB100 was a gift from Zsuzsanna Izsvak (Max Delbrück Center for Molecular Medicine, Berlin-Buch, Germany, plasmid 34879; http://n2t.net/addgene:34879; Addgene; RRID:Addgene_34879) ^104^. To create the RPE-Arf6-WT-eGFP cell line, an oligonucleotide encoding Arf6 fused to eGFP was generated by BioBasic using the ORF sequence of human Arf6, as per GenBank accession number NM_001663.4, followed by a spacer peptide sequence (5′-GGGGGGTCTGGTGGCAGTGGAGGGGGATCC-3′), followed by the ORF sequence of eGFP. This oligonucleotide sequence was subcloned into pSBtet-BP to generate pSBtet-BP-Arf6-eGFP. To create the RPE-Arf6-WT-UID cell line, an oligonucleotide encoding Arf6 fused to UltraID was generated by BioBasic using the ORF sequence of human Arf6 as per GenBank accession number NM_001663.4, followed by a spacer peptide sequence (5′-GGGGGGTCTGGTGGCAGTGGAGGGGGATCC-3′), followed by the ORF sequence of UltraID with a Myc tag as in ^79^. This oligonucleotide sequence was subcloned into pSBtet-BP to generate pSBtet-BP-Arf6-UltraID.

pSBtet-BP plasmids encoding Arf6-eGFP or Arf6-UltraID were co-transfected with pCMV(CAT)T7-SB100 into ARPE-19 cells using FuGENE HD transfection reagent as per the manufacturer’s protocol (Promega). Selection of stably engineered cells was performed in growth media supplemented with 2 μg/mL puromycin for 2-3 weeks. To induce the expression of Arf6-eGFP or Arf6-UltraID, cells were incubated with 1 μM doxycycline (dox; BioBasic) for 18-24 hr and stable expression was confirmed with western blotting and fluorescence microscopy. Experiments were performed in media supplemented with tetracycline-free fetal bovine serum (Gibco).

### Pharmacological treatments

All experiments began with cells incubated in low serum (0.1% FBS) culture media for 1 h before experimental treatments. Allosteric AMPK activation was induced with 100 μM A-769662 (Abcam) or an equal volume of DMSO (vehicle control; BioShop). Other pharmacological treatments were performed with 5 μM oligomycin (Cell Signaling Technology) or DMSO, 25 mM 2-deoxyglucose (2-DG; Millipore Sigma) or water, or 4 μM ikarugamycin (Millipore Sigma) or corresponding volume of vehicle (DMSO).

### Transient plasmid and siRNA transfections

Transient cDNA transfections were performed using Lipofectamine 2000 transfection reagent (Thermo Fisher Scientific) by a protocol adapted from the manufacturer’s protocol and as previously described ^66,101^. Cells were washed with 1X PBS and incubated in Opti-MEM (Gibco). For each well of a six-well plate, 2 μg of cDNA was precomplexed with 3 μL of Lipofectamine 2000 in Opti-MEM and incubated at room temperature for 15 min. DNA-reagent complexes were added dropwise onto adhered cells and incubated for 4 h. Following a 1X PBS wash, the transfection medium was replaced with regular growth media for 18-24 h before the start of the experiment.

siRNA transfections were performed using Lipofectamine RNAiMAX transfection reagent (Thermo Fisher Scientific) as per the manufacturer’s protocol and as previously described ^66,74,103^, using siRNA sequences as shown in **Table S1**. Cells were washed with 1X PBS and incubated in Opti-MEM (Gibco). Each siRNA was precomplexed with the transfection reagent in Opti-MEM, incubated for 15 min at room temperature, and added dropwise to adhered cells to a final siRNA concentration of 220 pmol/L. Cells were incubated with the siRNA complexes for 4 h, followed by a 1X PBS wash and replacement of growth media. Transfections were performed twice, 48 h and 24 h before each experiment.

### Immunofluorescence staining

Cell surface labelling of β1-integrin was performed as previously described ^19^. After the indicated treatments, cells were immediately placed on ice, washed with ice-cold 1X PBS supplemented with 1 mM MgCl_2_ and 1 mM CaCl_2_ (PBS++), and blocked with 3% BSA for 20 min. Cells were then incubated with either anti-β1-integrin antibody clone K20 (Novus Biologicals) or clone 4B7 (GeneTex) (**Table S2**) for 1 h on ice. Cells were then washed with 1X PBS++, fixed with 4% paraformaldehyde (PFA; Electron Microscopy Sciences) for 20 minutes, quenched with 100 mM glycine (BioShop), and incubated with the appropriate fluorescently conjugated secondary antibody (Jackson ImmunoResearch) (**Table S2**) for 1 h. Following extensive washing with 1X PBS++, coverslips were mounted in fluorescence mounting medium (Dako, Agilent).

For detection of specific proteins within CCPs and total cellular protein, the indicated treatments were performed and cells were immediately fixed in ice-cold 4% PFA for 20 min, quenched with 100 mM glycine, and permeabilized with 0.1% Triton X-100 (BioShop) for 10 min. Coverslips were blocked for one hour with 3% BSA, followed by incubation with the specific primary and fluorescently conjugated secondary antibodies (**Table S2**), each for 1 h. After extensive washing with 1X PBS, coverslips were either mounted in fluorescence mounting medium (Dako) or remained in 1X PBS at 4C until TIRF-M imaging.

Measurement of β1-integrin recycling to the plasma membrane was adapted from a similar assay used to measure recycling of the facilitative glucose transporter GLUT4 ^105^. Cells were simultaneously incubated with 100 μM A-769662 (or DMSO) and 1 mg/mL anti-β1-integrin K20 antibody (or not) for the indicated times (0-60 min) at 37C. Cells were then immediately placed on ice, washed with ice-cold 1X PBS++, fixed and permeabilized as above, and incubated with the appropriate fluorescently conjugated secondary antibody (Jackson ImmunoResearch) (**Table S2**) for 1 h. After extensive washing with PBS++, coverslips were mounted in fluorescence mounting medium (Dako). To measure β1-integrin recycling rate, each timed measurement of β1-integrin fluorescence intensity was divided by the fluorescence intensity of the total amount of β1-integrin detected in the cell, which was measured separately but in parallel in cells subjected to permeabilization with PBS supplemented with 0.1% TX-100 prior to labeling with anti-β1-integrin K20 antibody. The extent of recycling at each time point is thus expressed as the fluorescence intensity anti-β1-integrin recycling/total β1-integrin antibody fluorescence intensity.

To detect β1-integrin internalization into clathrin structures or EEA1-endosomes, cells were simultaneously incubated with 100 μM A-769662 (or DMSO) and 1 mg/mL β1-integrin antibody clone K20 (Novus Biologicals) for the indicated amount of time (2-5 min) at 37C. Cells were immediately placed on ice and the above protocol for detecting intracellular proteins was performed.

### Fluorescence microscopy

#### Widefield microscopy

Widefield epifluorescence microscopy experiments (**Figures 1B; 7G; 8E; S7D,G**) were performed on an Olympus IX83 Inverted Microscope with a 100×/1.4-NA objective coupled to a Hamamatsu ORCA-Flash 4.0 digital camera operating with cellSens software (Olympus Canada). Imaging was performed on samples mounted in fluorescence mounting medium (Dako) at room temperature.

#### TIRF and spinning disk confocal microscopy

TIRF-M experiments (**Figures 3A,C; 4C,E; 5; 6B,D; 7A,E; 8A,C; S2A-B; S3A-B,D; S4; S5C,F; S6A; S7C**) were performed on a Quorum Diskovery instrument comprised of a Leica DMi8 microscope equipped with a 63×/1.49-NA TIRF objective with a 1.8× camera relay (total magnification 108×). Imaging was done using 488, 561, or 647 nm laser illumination and 525/50, 620/60, and 700/75 nm emission filters and acquired using a Zyla 4.2Plus sCMOS camera (Andor). Image acquisition was performed in MetaMorph 7.10.3.279 (Molecular Devices). Fixed-cell TIRF-M imaging was done at room temperature with samples mounted in PBS. For live-cell imaging experiments, cells were maintained at 37C and 5% CO_2_ in phenol-free DMEM/F-12 media (Gibco) supplemented with 20 mM HEPES. For some TIRF-M experiments, widefield epifluorescence images were acquired of identical fields of view. For the experiments shown in **Figures 2B,D; S1; S6C,E; S7D**, the same instrument was operated in spinning disk confocal mode. For some experiments (**Figures 2D; S1**), each image series was obtained as a z-series with a step size of 200 nm; subsequent analysis was performed on the sum projections of these z-series.

For all microscopy images, final image processing was limited to linear adjustments of brightness/contrast, which were applied identically for all images of the same channel in an experiment, using ImageJ ^106^.

### Image Analysis

#### Analysis of widefield and spinning disk confocal fixed-cell images

Fluorescence intensity of cell surface β1-integrin was determined using ImageJ ^106^ after cell-surface labelling (non-permeabilized) and widefield or spinning disk fluorescence microscopy, as previously described ^19^. Briefly, a cell outline was determined manually and the mean fluorescence intensity corresponding to β1-integrin was measured in each cell. The nonspecific signal was determined using a similarly sized region of background (a region of the image not corresponding to any cell) and subtracted from the measured mean fluorescence intensity of β1- integrin for each cell. Cells not incubated with the primary antibody were used to verify the specificity of antibody labelling and detection. All measurements were subjected to statistical tests as described in the figure legends with a threshold of p < 0.05 for statistically significant differences between conditions. Data distribution was assumed to be normal but was not formally tested.

Colocalization analysis of β1-integrin within EEA1-endosomes was performed using the JaCoP plugin in ImageJ ^107^, as we have previously done in ^108^. Shown in **Figure 2D-E** and **Figure S1** are the Manders’ (M_1_) coefficients using β1-integrin signal as the primary channel and EEA1 signal as the secondary channel obtained using consistent thresholds across all image pairs. Cells not incubated with either primary antibody were used to verify the specificity of antibody labelling and detection. All measurements were subjected to statistical tests as described in the figure legends with a threshold of p < 0.05 for statistically significant differences between conditions. Data distribution was assumed to be normal but was not formally tested.

#### Analysis of fixed-cell TIRF images

Systematic, unbiased detection and analysis of clathrin labelled structures (CLSs) in fixed cells was done using custom software developed in Matlab (MathWorks Corp.), as previously described in ^33,72,74,109^. Briefly, diffraction-limited clathrin structures were detected using a Gaussian-based model method to approximate the point-spread function of eGFP-CLC or antibody-labelled clathrin structures (CLSs) in TIRF-M images (primary channel). The fluorescence intensity corresponding to various proteins in a secondary (or tertiary) channel (e.g., Arf6-eGFP, Dab2, AP2, epsin, or CALM) within CLSs was determined by the amplitude of the Gaussian model for the appropriate fluorescence channel for each CLS structure detected in the primary channel. As such, the measurements of fluorescently labeled proteins within CLSs represent their enrichment relative to the local background fluorescence in the immediate vicinity of the detected CLS. Similar measurements were done using widefield epifluorescence images (as the secondary channel) after CLS detection in the corresponding TIRF channel (as the primary channel).

Measurements (mean levels of various proteins within specified CLS subsets for each cell) were subjected to statistical tests as described in each figure legend, with a threshold of p < 0.05 for statistically significant differences between conditions. Data distribution was assumed to be normal but was not formally tested.

#### Identification of Arf6+ and Arf6-CLSs

To identify a subpopulation of clathrin structures containing Arf6-eGFP, we used a similar strategy as in ^66,74^, by establishing an arbitrary but systematic threshold of Arf6-eGFP intensity within CLS objects, that allowed classification of each object as Arf6+ or Arf6-. In this case, we used an Arf6-eGFP threshold of 0. Using this systematic threshold, we defined subsets of CLSs enriched in Arf6-eGFP as those with fluorescence intensity above this threshold.

#### Analysis of CCPs time-lapse image series

Automated detection, tracking, and analysis of CCPs was performed as previously described ^33,66,72,74^ following time-lapse TIRF-M imaging of RPE cells stably expressing eGFP-CLCa (eGFP-CLCa-RPE cells). Diffraction-limited clathrin structures were detected using a Gaussian-based model method to approximate the point-spread function and trajectories were determined from clathrin structure detections using u-track software ^110^. sCLSs were distinguished from bona fide CCPs based on quantitative and unbiased analysis of clathrin intensity progression in the early stages of structure formation, as previously described ^72^. Both sCLSs and CCPs represent nucleation events, but only bona fide CCPs represent structures that undergo stabilization, maturation, and in some cases scission to produce intracellular vesicles. We report the sCLS nucleation rate, CCP initiation rate, the fraction of CCPs that remain for < 15 seconds, and the ‘plateau intensity’ of eGFP-clathrin within these structures ^66^. The plateau intensity of eGFP-CLCa in TIRF or epifluorescence microscopy images as the mean fluorescence of that protein within each detected clathrin structure, measured within timepoints corresponding to 30% and 70% of the total lifetime of that structure, during which CCPs exhibit minimal growth or disassembly ^66^. Because CCPs are diffraction-limited objects, the amplitude of the Gaussian model of the fluorescence intensity of eGFP-CLCa informs about CCP size. All measurements were subjected to statistical tests as described in each figure legend with a threshold of p < 0.05 for statistically significant differences between conditions. Data distribution was assumed to be normal but was not formally tested.

### Biotin pulldown using Arf6-UltraID

Arf6-WT-UltraID cells were seeded onto 10 cm dishes and treated with 1 μM doxycycline 24 h prior to reaching 80-85% cell confluency. On the experiment day, culture medium was replaced with low serum (0.1% FBS) DMEM/F-12 for 1 hour, followed by incubation with 50 μM biotin (Thermo Fisher Scientific) and 100 μM A-769662 (or an equal volume of DMSO) for 15 minutes. Cells were placed on ice and extensively washed with 1X PBS prior to cell lysis with 1 mL Pierce RIPA buffer (Thermo Fisher Scientific) supplemented with 20LJnM Protease Inhibitor Cocktail and 1 mM DTT (both from BioShop). Lysates were passed through a 27.5-gauge syringe and rotated at 4C for 30 min. Samples were centrifuged at 14,000 × g for 10 min at 4C and the supernatant was incubated overnight with ∼50 μL of Dynabeads MyOne Streptavidin T1 bead slurry (Thermo Fisher Scientific) at 4C. The following day, beads were washed with lysis buffer (and inhibitors) three times. Proteins were eluted using a 1:1 ratio of bead volume:2X Laemmli sample buffer (LSB; 0.5 M Tris pH 6.8, 20% glycerol, 10% SDS; all from BioShop) and heated at 95C for 5 min. Input, unbound, and elution samples were collected at the appropriate steps and prepared for SDS-PAGE by adding 10% β-mercaptoethanol and 5% bromophenol blue (both from BioShop).

### Whole-cell lysates and western blotting

Cell lysis and western blotting were performed as previously described ^66,74,111^. After transfection or indicated treatments, cells were placed on ice and washed with 1X PBS. Whole cell lysates were prepared in 2X LSB with a protease and phosphatase inhibitor cocktail (1LJmM sodium orthovanadate, 10LJnM okadaic acid, and 20LJnM Protease Inhibitor Cocktail; all from BioShop). Lysates were supplemented with 10% β-mercaptoethanol and 5% bromophenol blue, heated at 65C for 15 min, and passed through a 27.5-gauge syringe. Proteins were resolved by glycine-Tris SDS–PAGE followed by transfer onto a 0.2 μm polyvinylidene fluoride membrane (Immobilon, Millipore). Membranes were blocked with 3% bovine serum albumin (BSA; BioShop) in TBST (Tris-buffered saline supplemented with Tween 20) and incubated with specific primary antibodies (**Table S2**) diluted in 1% BSA at 4C overnight. After TBST washes, membranes were incubated with the appropriate HRP-conjugated secondary antibodies (Cell Signaling Technology) (**Table S2**) at room temperature for 1 h. To detected biotinylated interactors of Arf6-WT-UltraID, membranes were blocked with 5% skim milk (BioShop) in TBST and incubated with Streptavidin-HRP (Cell Signaling Technology) at room temperature for 1 h. Following several TBST washes, bands were visualized using Luminata Crescendo HRP substrate (Millipore Sigma) on the Bio-Rad ChemiDoc Touch Imaging System. Western blot signals to detect the intensity corresponding to specific proteins (e.g., Arf6, AP2, ArfGAP3) were obtained by signal integration in an area corresponding to the appropriate lane and band for each condition. This measurement was normalized to the loading control (e.g., actin) signal. All measurements were subjected to statistical tests as described in the figure legends with a threshold of p < 0.05 for statistically significant differences between conditions. Data distribution was assumed to be normal but was not formally tested.

### Tfn and EGF ligand internalization assay (ELISA)

Measurement of the rate of internalization of Tfn and EGF ligands was done as previously described ^67^. Cells were simultaneously incubated with 100 μM A-769662 (or DMSO) and either 1 μg/mL biotin-xx-Tfn or 5 ng/mL biotin-xx-EGF (both from Life Technologies) for the indicated timepoints at 37C. Following ligand internalization, cells were immediately placed on ice and washed in ice-cold PBS++ to remove unbound ligand and arrest membrane traffic. Bound, uninternalized ligands were subsequently quenched by sequential incubation with free avidin (BioBasic) (28.6 μg/mL for Tfn and 5 μg/ml for EGF) for 30 min on ice, followed by biocytin (Santa Cruz Biotechnology) (27.3 μg/ml for Tfn and 7.69 μg/ml for EGF) for 15 min on ice. Cells were then washed with ice-cold PBS++ and solubilized by incubation in Superblock (Thermo Fisher Scientific) supplemented with 0.01% Triton X-100 and 0.01% SDS. Cell lysates were plated onto enzyme-linked immunosorbent assay (ELISA) plates coated with either anti-Tfn (Bethyl Laboratories) or anti-EGF (Millipore Sigma) antibodies, followed by detection of biotin-xx-Tfn or biotin-xx-EGF (free, unbound to avidin) by incubation with streptavidin conjugated to horseradish peroxidase (POD; BioBasic), and detection of bound POD by o-phenylenediamine assay. Absorbance readings corresponding to internalized biotin-xx-Tfn or biotin-xx-EGF were then normalized to the total levels of surface-ligand binding measured at 4C and not subjected to avidin-biocytin quenching, performed in parallel for each condition. All measurements were subjected to statistical tests as described in each figure legend, with a threshold of p < 0.05 for statistically significant differences between conditions. Data distribution was assumed to be normal but was not formally tested.

### BioID

BioID of Arf6-WT-BirA*, Arf6-Q67L-BirA*, Arf6-T44N-BirA*, and Arf6-TA-BirA* were performed as previously described ^112^. HeLa cells were cultured to approximately ∼75% confluency, and bait expression along with biotin labeling were induced simultaneously by adding 1 µg/ml tetracycline and 50 µM biotin for 24 h. After induction, cells were rinsed, scraped into 1 mL of PBS, collected by centrifugation at 500LJ×LJ*g* for 3 min, and stored at −80C until further processing. For lysis, frozen cell pellets were thawed on ice and an ice-cold lysis buffer (4:1 (v/w) ratio) was added (50 mM Tris-HCl, pH 7.5, 150 mM NaCl, 1% Nonidet P-40 substitute (IGEPAL-630), 0.4% SDS, 1.5 mM MgCl2, 1 mM EGTA, benzonase, protease inhibitors). Cells were resuspended using a P1000 pipette tip (∼10–15 aspirations) and subjected to a rapid freeze/thaw cycle (dry ice to 37C water bath). Lysates were rotated at 4C for 30 min and centrifuged at 16,000 × g for 20 min at 4C. Supernatants were collected and incubated overnight at 4C with 20 µL (packed beads) of pre-washed streptavidin-Sepharose (GE) while rotating. The beads were collected by centrifugation at 500LJ×LJ*g* for 2 min, the supernatant was discarded, and the beads were transferred to new tubes with 500 µL of lysis buffer. The beads were washed once with SDS wash buffer (50 mM Tris-HCl, pH 7.5, 2% SDS), twice with lysis buffer, and three times with 50 mM ammonium bicarbonate (ABC), pH 8.0. Between each wash, beads were centrifuged at 500LJ×LJ*g* for 30LJsec. For digestion, beads were resuspended in 100 µL of ABC containing 1 µg of sequencing grade trypsin and gently mixed for 4 h at 37C. A fresh 1 µg of trypsin was added and digestion continued overnight. The supernatant was collected by centrifugation at 500LJ×LJ*g* for 2 min, and the beads were washed with 100 µL of molecular biology grade H_2_O and pooled with peptides. Digestion was stopped by acidification with formic acid (2% final concentration) and peptides were dried by vacuum centrifugation.

### Mass spectrometry acquisition using TripleTOF mass spectrometers

Each sample (5 μL in 2% formic acid; equivalent to 1/8th of a 15 cm tissue culture dish) was directly loaded on to an equilibrated HPLC column at a flow rate of 800LJnL/min. Peptide elution occurred over a 90 min gradient using an Eksigent ekspert™ nanoLC 425 nano-pump (Eksigent, Dublin CA), followed by analysis on a TripleTOF^TM^ 6600 instrument (AB SCIEX, Concord, Ontario, Canada). The gradient was delivered at 400LJnL/min, starting from 2% acetonitrile with 0.1% formic acid to 35% acetonitrile with 0.1% formic acid over 90 min. This was followed by a 15 min clean-up at 80% acetonitrile with 0.1% formic acid, and a 15 min equilibration period back to 2% acetonitrile with 0.1% formic acid, for a total run time of 120 min. To minimize carryover between samples, the analytical column was washed for 2 h using an alternating sawtooth gradient from 35% to 80% acetonitrile with 0.1% formic acid at a flow rate of 1500nL/min, holding each gradient concentration for 5 min. Analytical column and instrument performance were verified after each sample by loading 30LJfmol bovine serum albumin (BSA) tryptic peptide standard along with 60LJfmol α-casein tryptic digest and running a short 30 min gradient. TOF MS mass calibration was performed on BSA reference ions to adjust for mass drift and verify peak intensity before running the next sample. Samples were analyzed in data-dependent acquisition (DDA) mode, with each cycle consisting of one 250 ms MS1 TOF survey scan (400-1800LJDa), followed by ten 100 ms MS2 candidate ion scans (100- 1800LJDa) in high sensitivity mode. Only ions with a charge of 2+ to 5+ that exceeded a threshold of 300LJcps were selected for MS2, and former precursors were excluded for 7 s after one occurrence.

### SAINT analysis and data visualization

SAINTexpress (version 3.6.1 (45)) was used to score proximity interactions from DDA data. This tool estimates the probability of a true proximity interaction for each prey protein identified with a given bait, relative to negative control runs, using spectral counting as a proxy for protein abundance. Bait runs (two biological replicates each) were compared against twelve negative control runs consisting of four BirA*-FLAG-only samples, four 3xFLAG-only samples, and four EGFP-BirA*-FLAG samples. These were consolidated into four “virtual controls” that maximize scoring stringency, simulating the worst-case scenario where proteins are detected either because they are endogenously biotinylated in the absence of BirA*, or frequently biotinylated by recombinant BirA* expression. Prey proteins with a false discovery rate (FDR)LJ<LJ1% (Bayesian estimation based on the distribution of the Averaged SAINT scores across both biological replicates) were considered high-confidence proximity interactions. Dotplots were generated using ProHits-viz (prohits-viz.org) ^113^, which retrieves the quantitative values for all baits once a prey protein passes the selected FDR threshold (1%) (**Table S3**).

## Notes

### Competing Interest Statement

The authors have declared no competing interest.

