## Supplemental Materials for "Cargo-selective regulation of clathrin-mediated endocytosis by AMP-activated protein kinase"

**Figure S1**: AMPK activation promotes β1-integrin accumulation in early endosomes.
**Figure S2**: A-769662 treatment reduces the levels of eGFP-clathrin within each CCP.
**Figure S3**: AMPK activation promotes β1-integrin recruitment to clathrin structures in a cargo-selective manner.
**Figure S4**: AMPK activation enhances β1-integrin abundance in clathrin structures in a Dab2-dependent manner.
**Figure S5**: Extended BioID dataset and clathrin-coated structure analysis for Arf6 GTP-binding mutants.
**Figure S6**: AMPK regulates Arf6 in clathrin structures in an ArfGAP3 dependent-manner, and validation experiments for Arf6-UltraID.
**Figure S7**: Arf6 and ArfGAP3 are required to enhance Dab2 in clathrin structures and reduce cell surface β1-integrin in response to AMPK activation.
**Figure S8**: Arf6 silencing has modest effects on CCP initiation, dynamics, and size.
**Table S1**: siRNA sequences used in this study
**Table S2**: Antibodies for western blotting and immunofluorescence

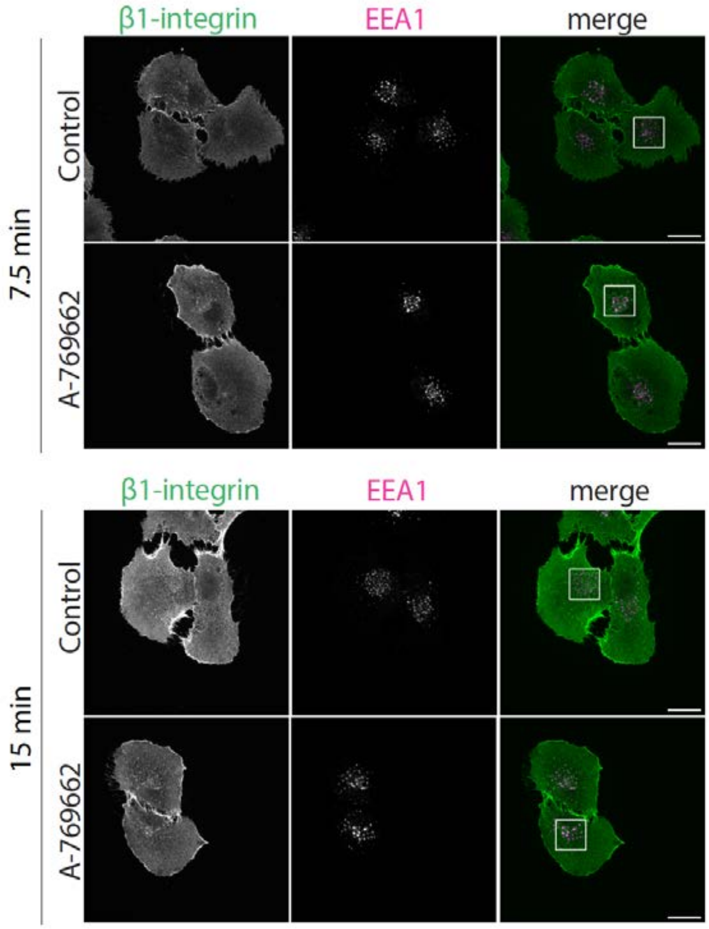


**Figure S1. AMPK activation promotes β1-integrin accumulation in early endosomes.**
ARPE-19 cells simultaneously treated with 1 mg/mL anti-β1-integrin (antibody clone K20) and either 100 μM A-769662 or vehicle control for 15 minutes. Shown are full field of view z-series slices of spinning disk confocal images of fixed cells labelled for β1-integrin and early endosome marker EEA1. The white square indicates cropped region displayed in **Figure 2D**. Scale bar represents 20 μm.

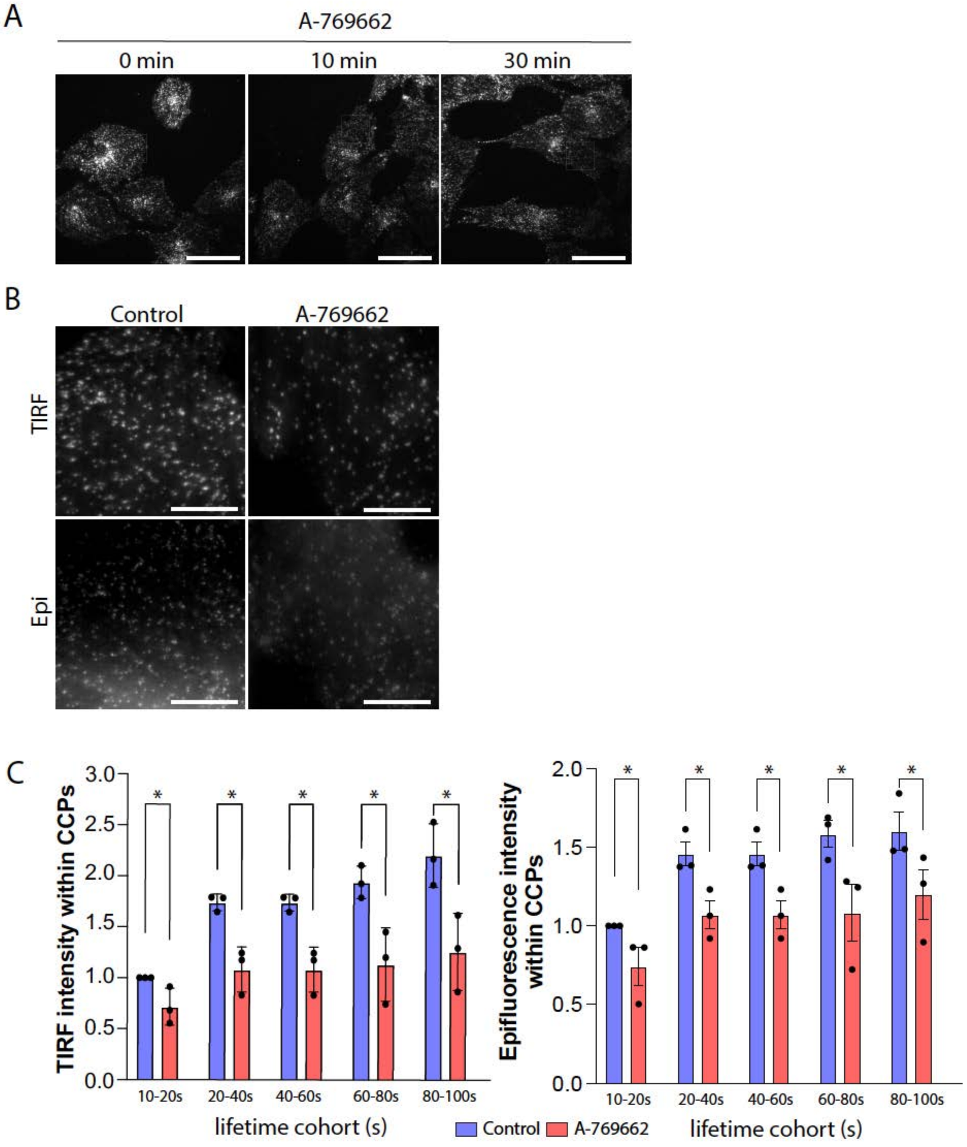


**Figure S2. A-769662 treatment reduces the levels of eGFP-clathrin within each CCP.**
(A) Representative time-lapse TIRF-M images of eGFP-CLCa-RPE cells treated with 100 µM A769662 or vehicle control for the indicated times. Scale bar represents 40 µm. (B) Representative TIRF-M and widefield epifluorescence images of clathrin structures acquired concomitantly after treatment with 100 µM A-769662 or vehicle control for 10 minutes. Scale bar represents 5 µm. Shown is the mean ± SD of eGFP-clathrin fluorescence intensity in the TIRF (*left*) and epifluorescence (*right*) channels within each lifetime cohort determined by automated detection, tracking, and analysis of clathrin structures from three independent experiments as described in *Methods*. Control: k (cells) = 19 and n (CLSs) = 12112; A-769662: k (cells) = 21 and n (CLSs) = 5686; * $p<0.05$ [two-way ANOVA with Šídák's post-hoc test].

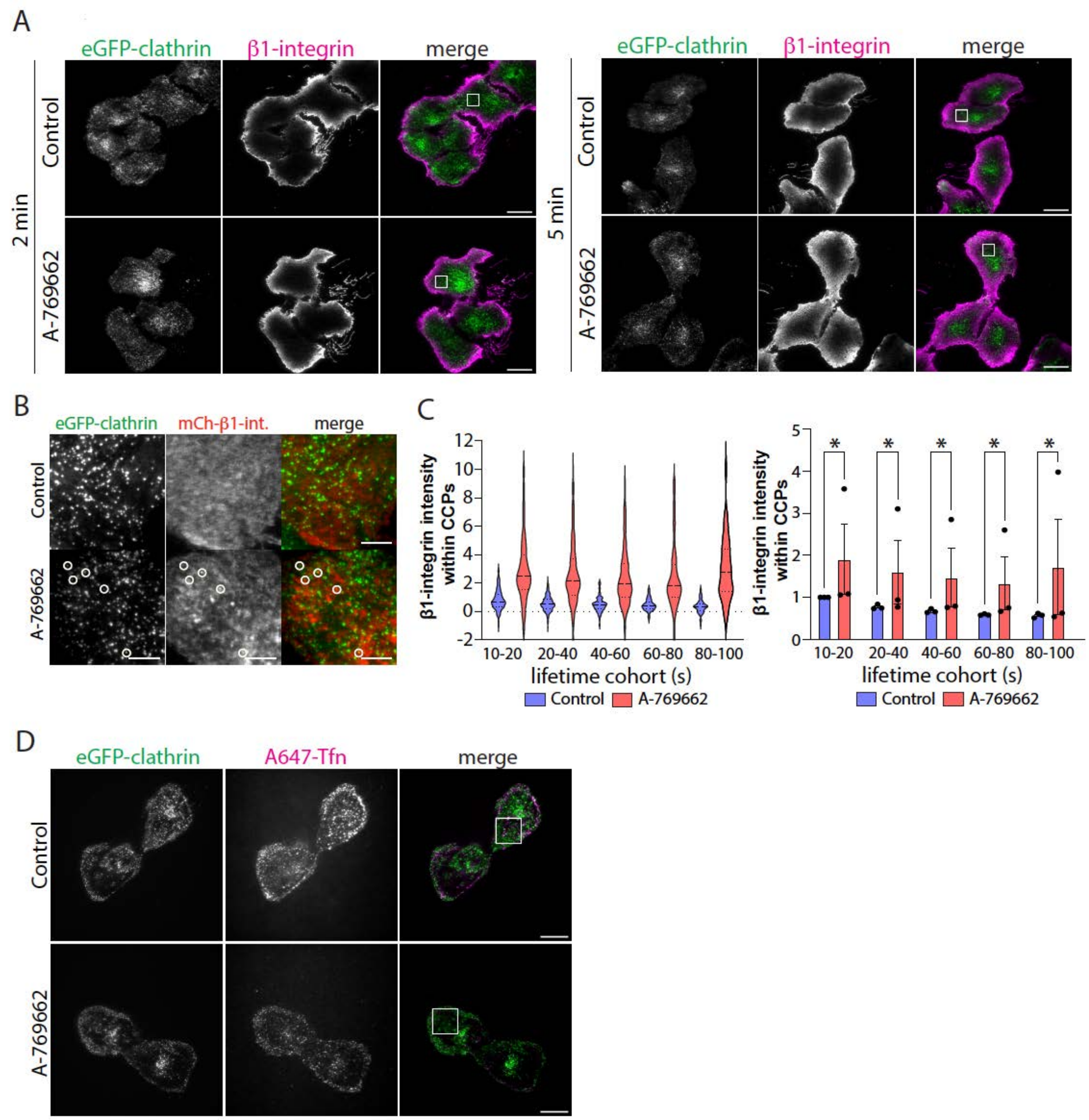


**Figure S3. AMPK activation promotes β1-integrin recruitment to clathrin structures in a cargo-selective manner.**
(A) eGFP-CLCa-RPE cells simultaneously treated with 1 mg/mL anti-β1-integrin (antibody clone K20) and either 100 μM A-769662 or vehicle control were fixed at the indicated timepoints and imaged by TIRF-M. The white square indicates the cropped region displayed in **Figure 4C**. Scale bar represents 20 μm. (B) eGFP-CLCa-RPE cells transiently transfected with mCherry-β1-integrin were treated with 100 μM A-769662 or vehicle control for 15 minutes. Scale bar represents 5 μm. Time-lapse TIRF-M image series were subject to automated detection, tracking, and analysis of clathrin structures and β1-integrin as described in *Methods*. (C) Shown is a representative experiment of β1-integrin fluorescence intensity detected in CCPs within lifetime cohorts (*left*) and the β1-integrin fluorescence intensity means ± SD from three independent experiments (*right*). Control: k (cells) = 11 and n (CLSs) = 11162; A-769662: k (cells) = 15 and n (CLSs) = 7886. * $p<0.05$ [two-way ANOVA with Šídák's post-hoc test] (D) eGFP-CLCa-RPE cells were treated simultaneously with A647-Tfn and 100 μM A-769662 or vehicle control treatment for 5 minutes, fixed, and imaged by TIRF-M. The white square indicates the cropped region displayed in **Figure 4E**. Scale bar represents 20 μm.

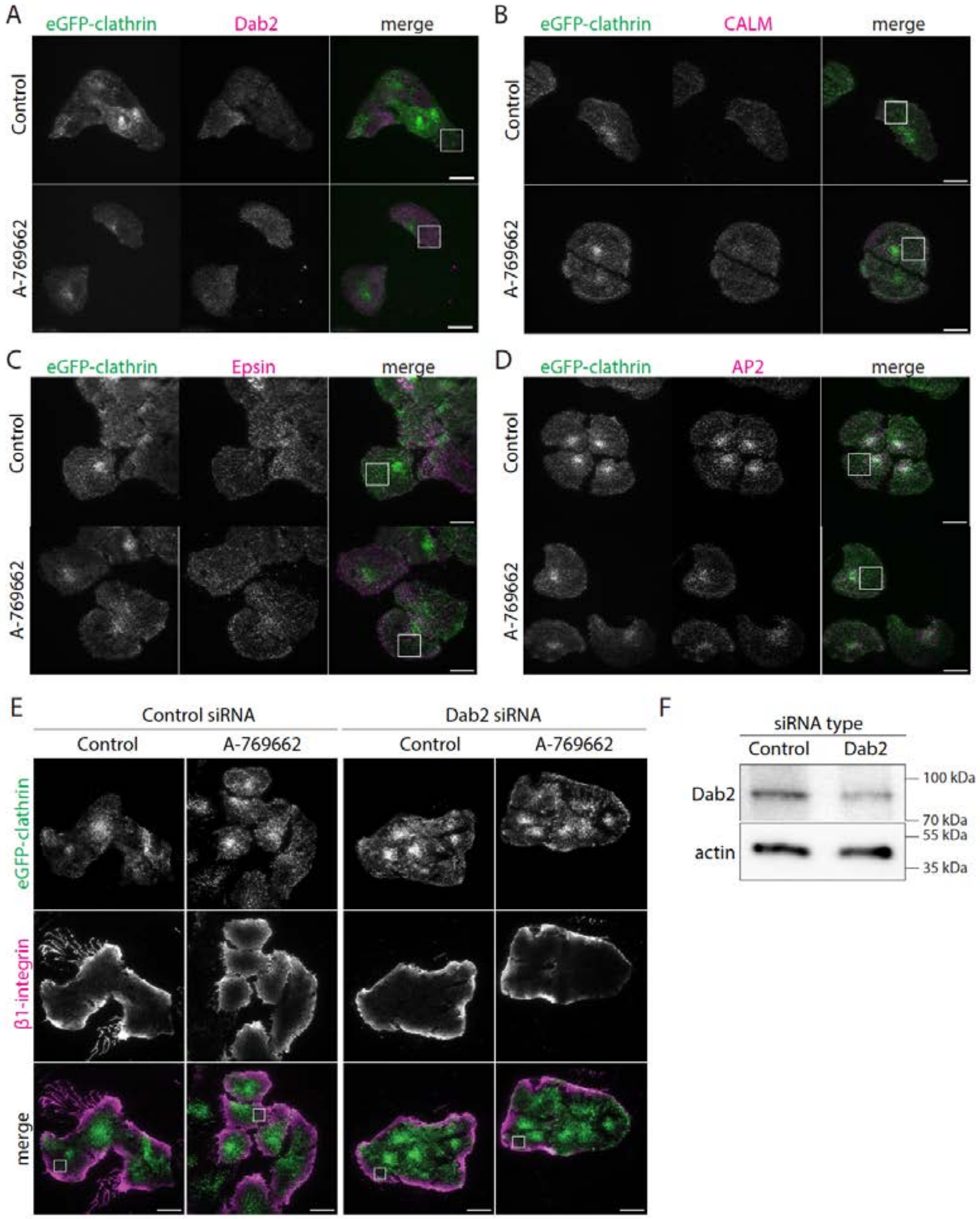


**Figure S4. AMPK activation enhances β1-integrin abundance in clathrin structures in a Dab2-dependent manner.**

eGFP-CLCa-RPE cells treated with 100 μM A-769662 or vehicle control for 5 minutes. Fixed cells were labelled with antibodies against (A) Dab2, (B) CALM, (C) epsin, or (D) AP2 and imaged by TIRF-M. The white square represents the cropped field of view shown in **Figure 5A,C,G,E**. Scale bars represent 20 μm. (E) eGFP-CLCa-RPE cells transfected with Dab2 siRNA or non-targeting control siRNA were simultaneously treated with 1 mg/mL anti-β1-integrin (antibody clone K20) and either 100 μM A-769662 or vehicle control at 37C for 5 minutes. Cells were fixed at the and imaged by TIRF-M. The white square represents the cropped field of view shown in **Figure 5I**. Scale bar represents 20 μm. (F) Representative immunoblot of eGFP-CLCa-RPE cells transfected with Dab2 siRNA or non-targeting control siRNA.

A

| PANTHER Protein Class | Client Text Box Input (504) | Client Text Box Input (expected) | Client Text Box Input (fold Enrichment) | Client Text Box Input (raw P-value) | Client Text Box Input (FDR) |
|---|---|---|---|---|---|
| SNARE protein (PC00034) | 10 | 0.76 | 13.17 | 1.98E-09 | 6.48E-08 |
| vesicle coat protein (PC00235) | 5 | 1.1 | 4.54 | 4.71E-03 | 3.84E-02 |
| membrane traffic protein (PC00150) | 37 | 10.6 | 3.49 | 4.68E-11 | 2.29E-09 |
| small GTPase (PC00208) | 16 | 3.06 | 5.23 | 7.07E-08 | 1.26E-06 |
| integrin (PC00126) | 4 | 0.69 | 5.83 | 4.56E-03 | 3.89E-02 |
| amino acid transporter (PC00046) | 4 | 0.64 | 6.28 | 3.46E-03 | 3.09E-02 |
| secondary carrier transporter (PC00258) | 25 | 5.85 | 4.27 | 1.16E-09 | 4.55E-08 |
| transporter (PC00227) | 68 | 25.32 | 2.69 | 9.71E-14 | 6.35E-12 |
| primary active transporter (PC00068) | 18 | 5.53 | 3.25 | 1.25E-05 | 1.89E-04 |
| ligase (PC00142) | 6 | 1.32 | 4.54 | 2.00E-03 | 1.96E-02 |
| chaperonin (PC00073) | 4 | 0.32 | 12.56 | 2.13E-04 | 2.79E-03 |
| G-protein (PC00020) | 21 | 4.68 | 4.49 | 1.03E-08 | 2.89E-07 |
| non-motor actin binding protein (PC00165) | 8 | 2.13 | 3.75 | 1.32E-03 | 1.43E-02 |
| non-receptor serine/threonine protein kinase (PC00167) | 17 | 8.42 | 2.02 | 6.88E-03 | 5.00E-02 |
| protein-binding activity modulator (PC00095) | 35 | 19.59 | 1.79 | 9.57E-04 | 1.10E-02 |
| translation factor (PC00223) | 8 | 2.77 | 2.89 | 6.66E-03 | 5.02E-02 |
| translational protein (PC00263) | 18 | 8.2 | 2.19 | 1.87E-03 | 1.93E-02 |

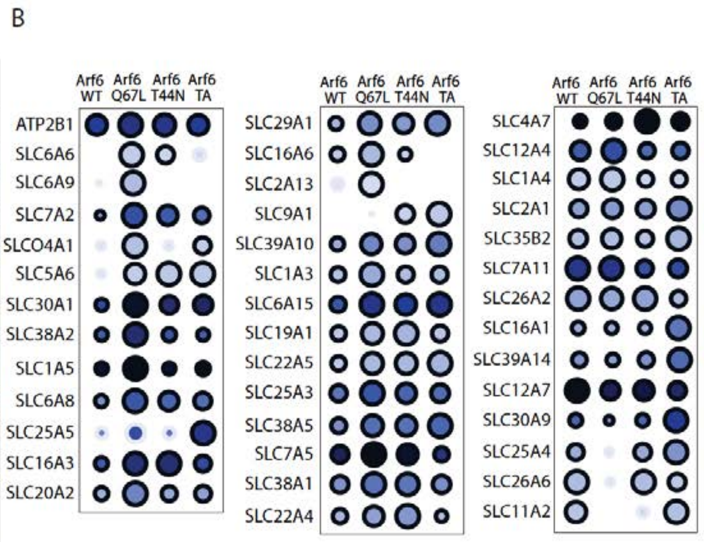


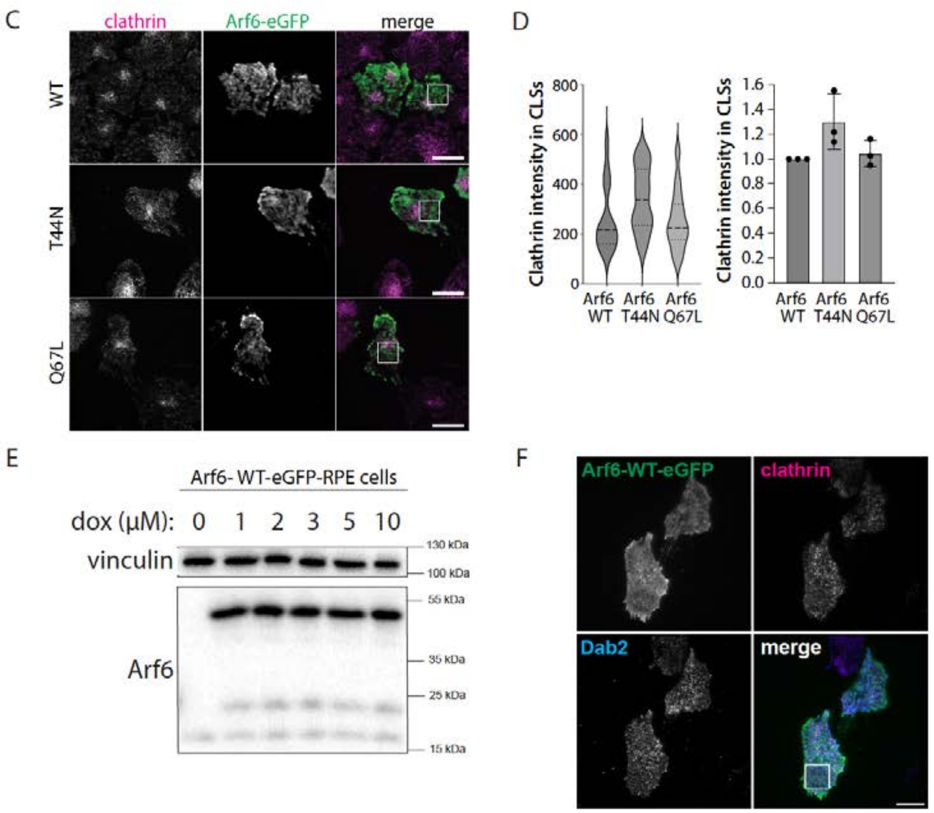


**Figure S5. Extended BioID dataset and clathrin-coated structure analysis for Arf6 GTP-binding mutants.**

(A) PANTHER protein class analysis of proteins detected within the interactome of Arf6 constructs. (B) Dot plot representing the extended BioID dataset of prey proteins identified with Arf6-WT-BirA*, Arf6-Q67L-BirA* (GTP-bound), or Arf6-T44N-BirA* (GDP-bound), or Arf6-TA-BirA* (fast-cycling mutant, T157A) enriched over endogenous biotinylation (untransfected) and non-specific pan-cellular biotinylation

(BirA* alone—BFDR ≤ 1%, SAINT). (C) RFP-CLCa-RPE cells transiently transfected Arf6-WT-eGFP, Arf6-Q67L-eGFP, or Arf6-T44N-eGFP were visualized by TIRF-M. The white square represents the cropped field of view shown in **Figure 6B**. Scale bar represents 20 μm. (D) Automated detection and analysis of the fluorescence intensity of RFP-clathrin puncta was performed as described in *Methods*. The fluorescence intensity of clathrin structures is shown as a representative experiment (*left*) and as the mean ± SD from three independent experiments (*right*). The number of cells and CLSs analyzed in each experiment are as described in **Figure 6C**. *$p<0.05$ [one-way ANOVA with Dunnett's post-hoc test]. (E) Arf6-WT-eGFP-RPE cells were stimulated with the indicated concentrations of doxycycline for 24 hours. Shown is a representative immunoblot of whole cell lysates probed with anti-Arf6 and anti-vinculin antibodies. Arf6-WT-eGFP is detected at ~50 kDa; endogenous Arf6 is detected at ~19 kDa. (F) Arf6-WT-eGFP-RPE cells treated with 1 μM doxycycline for 24 hours were fixed, labelled with anti-Dab2 and anti-clathrin antibodies, and imaged by TIRF-M. The white square represents the cropped field of view shown in **Figure 6D**. Scale bar represents 20 μm.

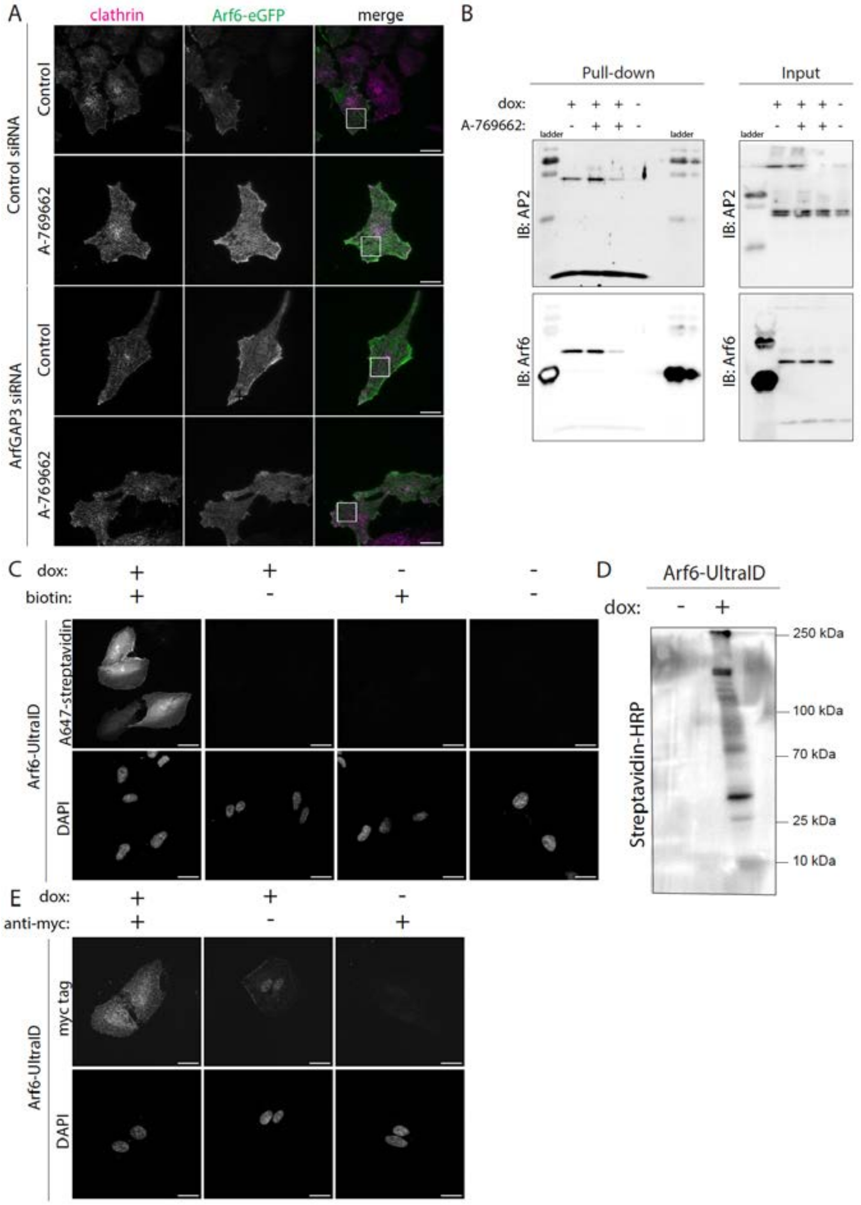


**Figure S6. AMPK regulates Arf6 in clathrin structures in an ArfGAP3 dependent-manner, and validation experiments for Arf6-UltraID.**
(A) Arf6-WT-eGFP-RPE cells transfected with ArfGAP3 siRNA or non-targeting control siRNA were stimulated with 1 μM doxycycline for 24 hours, followed by treatment with 100 μM A-769662 or vehicle control for 5 minutes. Shown are TIRF-M images of fixed cells labelled with an anti-clathrin antibody. The white square represents the cropped field of view shown in **Figure 7A** and **Figure 8A**. Scale bar represents 20 μm. (B) Arf6-WT-UltraID cells stimulated with 1 μM doxycycline (dox) for 24 hours were treated with 50 μM biotin in the presence of 100 μM A-769662 or vehicle control for 15 minutes. Shown is the uncropped representative immunoblot of whole cell lysates (input) and biotinylated interactors isolated by streptavidin pulldown (IP) shown in **Figure 7C**. (C) Arf6-WT-UltraID cells were stimulated with 1 μM doxycycline for 24 hours and treated with 50 μM biotin for 15 minutes. Biotinylated interactors were visualized with Alexa Fluor 647 Streptavidin on a spinning disk confocal microscope. Scale bar represents 20 μm. (D) Immunoblot of Arf6-WT-UltraID cells stimulated with 1 μM doxycycline for 24 hours and

treated with 50 μM biotin for 15 minutes. Biotinylated interactors were visualized from whole cell lysate probed with streptavidin-HRP. (E) Arf6-WT-UltraID cells were stimulated with 1 μM doxycycline for 24 hours, fixed, and labelled with an anti-myc tag antibody. Shown are representative spinning disk confocal images. Scale bar represents 20 μm.

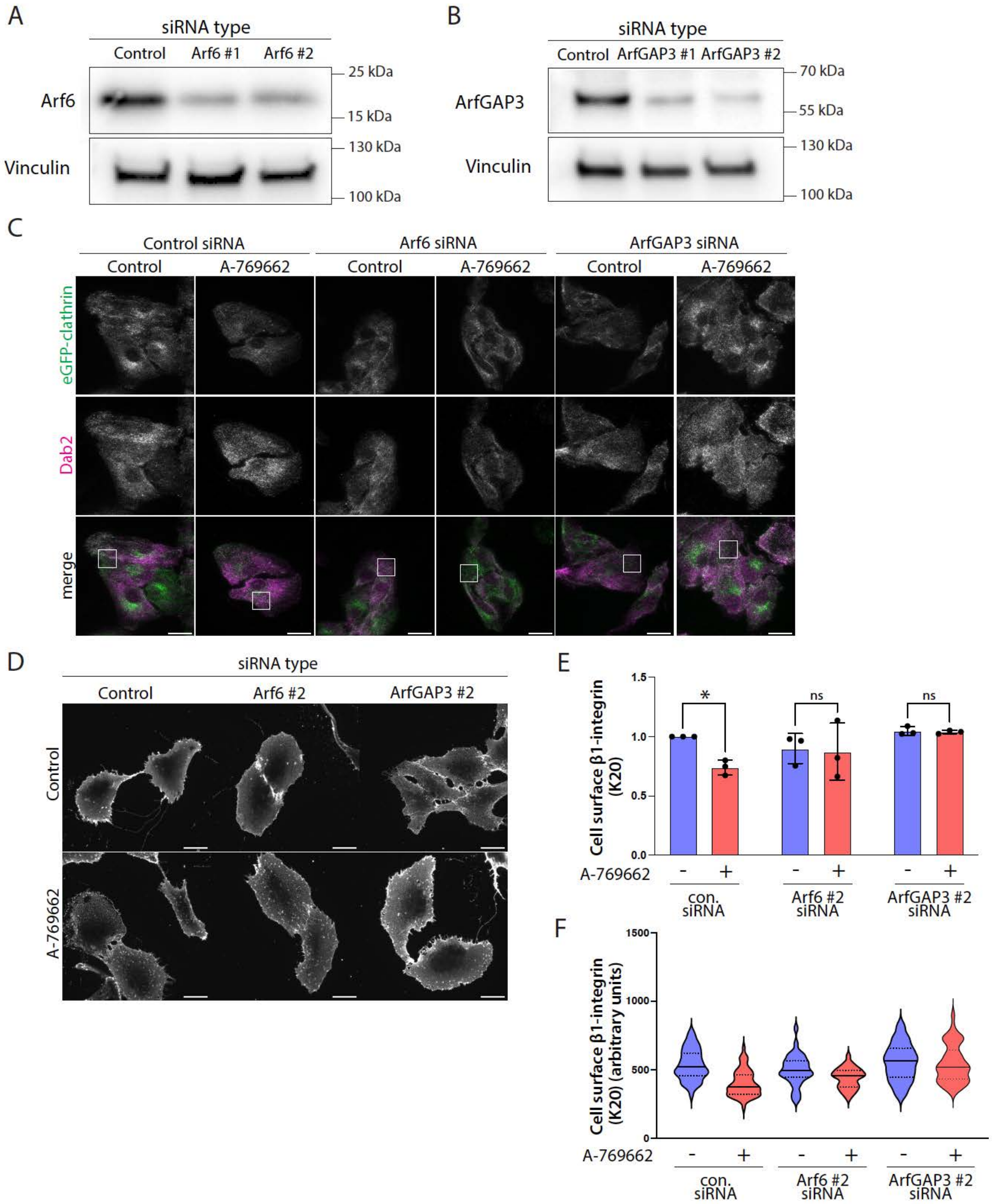

A
siRNA type
Control
Arf6 #1
Arf6 #2
Arf6
Vinculin
25 kDa
15 kDa
130 kDa
100 kDa
B
siRNA type
Control
ArfGAP3 #1
ArfGAP3 #2
ArfGAP3
Vinculin
70 kDa
55 kDa
130 kDa
100 kDa
C
Control siRNA
Arf6 siRNA
ArfGAP3 siRNA
Control
A-769662
eGFP-clathrin
Dab2
merge
D
siRNA type
Control
Arf6 #2
ArfGAP3 #2
Control
A-769662
E
Cell surface β1-integrin (K20)
1.5
1.0
0.5
0.0
*
ns
ns
A-769662
-
+
con. siRNA
Arf6 #2 siRNA
ArfGAP3 #2 siRNA
F
Cell surface β1-integrin (K20) (arbitrary units)
1500
1000
500
0
A-769662
-
+
con. siRNA
Arf6 #2 siRNA
ArfGAP3 #2 siRNA

G

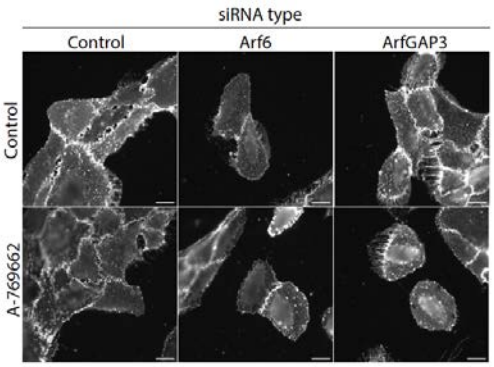


H

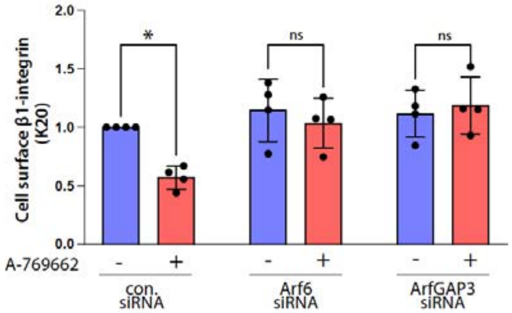


I

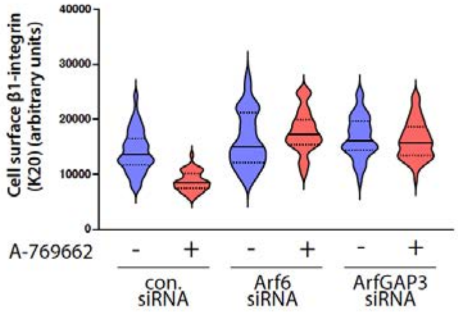


**Figure S7. Arf6 and ArfGAP3 are required to enhance Dab2 in clathrin structures and reduce cell surface β1-integrin in response to AMPK activation.**
(A) Representative immunoblot of eCLCa-RPE cells transfected with Arf6 siRNA sequence #1, Arf6 siRNA sequence #2, or non-targeting control siRNA. (B) Representative immunoblot of eGFP-CLCa-RPE cells transfected with ArfGAP3 siRNA sequence #1, ArfGAP3 siRNA sequence #2, or non-targeting control siRNA. (C) eGFP-CLCa-RPE cells transfected with Arf6 siRNA, ArfGAP3 siRNA, or non-targeting control siRNA were treated with 100 μM A-769662 or vehicle control for 5 minutes and labelled with an anti-Dab2 antibody. The white square represents the cropped field of view shown in **Figure 7E** and **Figure 8C**. Scale bar represents 20 μm. (D) ARPE-19 cells transfected with Arf6 siRNA #2, ArfGAP3 siRNA #2, or non-targeting control siRNA were treated with 100 μM A-769662 or vehicle control for 5 minutes and antibody labelled for cell surface β1-integrin (antibody clone K20). Scale bar represents 20 μm. Mean fluorescence intensity of cell surface β1-integrin is shown as (E) a representative experiment and as (F) the mean ± SD from three independent experiments that analyzed 30-50 individual cells per condition. * $p<0.05$ [two-way ANOVA with a Šídák post-hoc test]. (G) ARPE-19 cells transfected with Arf6 siRNA (#1), ArfGAP3 siRNA (#1), or non-targeting control siRNA were treated with 100 μM A-769662 or vehicle control for 5 minutes and antibody labelled for cell surface β1-integrin (antibody clone K20). Scale bar represents 20 μm. Mean fluorescence intensity of cell surface β1-integrin is shown as (H) a representative experiment and as (I) the mean ± SD from three independent experiments that analyzed 30-50 individual cells per condition. * $p<0.05$ [two-way ANOVA with a Šídák post-hoc test].

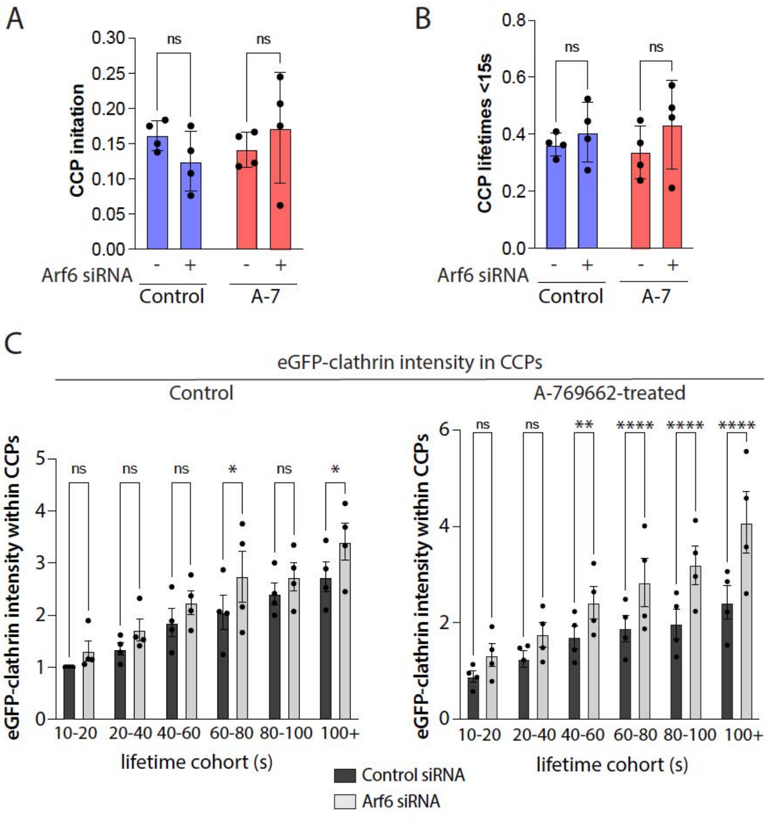


**Figure S8. Arf6 silencing has modest effects on CCP initiation, dynamics, and size.**
eGFP-CLCa-RPE cells treated with siRNA targeting Arf6 or control (non-targeting) siRNA. Cells were subjected to time-lapse TIRF-M imaging while being treated with 100 µM A-769662 (or not). Time-lapse image series were analyzed by automated detection and tracking of CLSs as described in *Methods* to identify bona fide CCPs. Shown is the mean ± SD from four independent experiments for (A) CCP initiation, (B) CCPs with lifetimes <15 seconds, and (C) eGFP-clathrin intensity within CCPs. Control siRNA, control: k (cells) = 32 and n (CLSs) = 4975; control siRNA, A-769662: k (cells) = 32 and n (CLSs) = 2153; Arf6 siRNA, control: k (cells) = 27 and n (CLSs) = 3717; Arf6 siRNA, A-769662: k (cells) = 23 and n (CLSs) = 3182. * $p<0.05$ [two-way ANOVA with a Šídák post-hoc test].

**Table S1. siRNA sequences used in this study**

| Target | Sense | Anti-sense |
|---|---|---|
| Dab2 | GUU CAA AGG CGA UGG UGU AUU | UAC ACC AUC GCC UUU GAA CUU |
| Arf6 (#1) | ACG UGG AGA CGG UGA CUU AUU | UAA GUC ACC GUC UCC ACG UUU |
| Arf6 (#2) | CAA CAA UCC UGU ACA AGU UUU | AAC UUG UAC AGG AUU GUU GUU |
| ArfGAP3 (#1) | GAU GAA CAU UAG UGG CAA AUU | UUU GCC ACU AAU GUU CAU CUU |
| ArfGAP3 (#2) | GAA ACC AAA UCA AGC UAA AUU | UUU AGC UUG AUU UGG UUU CUU |
| Control siRNA (Con7) | CGU ACU GCU UGC GAU ACG GUU | CCG UAU CGC AAG CAG UAC GUU |

**Table S2. Antibodies for western blotting and immunofluorescence**

| | Company | Product Code |
|---|---|---|
| **Immunofluorescence Antibodies** | | |
| Integrin beta 1/CD29 (4B7) | GeneTex | GTX20035 |
| Integrin beta 1/CD29 (K20) | Novus Biologicals | NBP2-52665 |
| EEA1 (E9Q6G) | Cell Signaling Technology | 48453S |
| Dab2 (D7O9T) | Cell Signaling Technology | 12906S |
| CALM (PICALM) | Sigma Aldrich | HPA019061 |
| Epsin | Abcam | ab75879 |
| AP2 ($\mu$2) (AP6) | Abcam | ab2730 |
| Clathrin heavy chain (X22) | Generated in-house from [112] | |
| Myc-tag | Cell Signaling Technology | 2272S |
| Alexa Fluor 647 Streptavidin | Jackson ImmunoResearch | 016-600-084 |
| Donkey anti-rabbit, donkey anti-mouse (Cy5, Cy3, 488) | Jackson ImmunoResearch | |
| **Western Blotting Antibodies** | | |
| pACC | Cell Signaling Technology | 3661S |
| Total ACC | Cell Signaling Technology | C83B10 |
| Pan-Actin (D18C11) | Cell Signaling Technology | 8456S |
| AP2 ($\mu$2) (AP50) | BD Biosciences | 611351 |
| Streptavidin-HRP | Cell Signaling Technology | 3999S |
| Arf6 (D12G10) | Cell Signaling Technology | 5740S |
| ArfGAP3 | Sigma Aldrich | HPA000638 |
| Vinculin (7F9) | Santa Cruz Biotechnology | sc-73614 |
| Anti-mouse IgG, anti-rabbit IgG (HRP-linked) | Cell Signaling Technology | 7076S, 7074S |
